# Twinfilin-1 phosphorylation in reelin signaling regulates actin dynamics and spine development

**DOI:** 10.1101/2025.06.18.660483

**Authors:** Geyao Dong, Daisuke Mori, Tetsuo Matsuzaki, Rinako Tanaka, Norimichi Itoh, Takaaki Matsui, Ayato Sato, Yuko Arioka, Hiroki Okumura, Ryota Fukaya, Hiroshi Kuba, Taku Nagai, Toshitaka Nabeshima, Hiroaki Ikesue, Takao Kohno, Mitsuharu Hattori, Kozo Kaibuchi, Norio Ozaki, Hiroyuki Mizoguchi, Kiyofumi Yamada

**Author notes:** Corresponding author. (H. Mizoguchi). Corresponding author. *E-mail addresses:* (K. Yamada).

## Abstract

Reelin is an extracellular glycoprotein essential for neuronal migration, spine development, and synaptic plasticity. Impaired reelin signaling is linked to neurological disorders, including schizophrenia and autism. While reelin mutant (*reeler*) mice exhibit behavioral deficits associated with impaired spine formation, the underlying molecular mechanisms remain unclear. We identified Twinfilin-1 (Twf1) as a downstream effector of reelin signaling via phosphoproteomic analysis, based on its reduced tyrosine phosphorylation in *reeler* mice. We found that Src regulated Twf1 phosphorylation at tyrosine 309, and reelin stimulation increased Twf1 phosphorylation in neurons, an effect blocked by the Src inhibitor PP2. A phospho-resistant Twf1 mutant (Twf1 Y309F) showed reduced capping protein binding and a lower F/G-actin ratio. *Twf1^Y309F^* mice exhibited cognitive deficits, reduced spine density, smaller spine head size, and a decreased F/G-actin ratio in synaptosomes. These findings highlight Twf1 phosphorylation as a key component of reelin signaling involved in actin remodeling and spine development.

## 1. Introduction

Reelin is an extracellular glycoprotein produced by Cajal-Retzius cells in the marginal zone of the developing brain, where it regulates neuronal migration and positioning [1,2]. In the adult brain, reelin is expressed by γ-aminobutyric acid (GABA)-ergic interneurons and plays a crucial role in spine development and synaptic plasticity [3–5]. Human genetic studies have identified rare variants of the *RELN* gene as a genetic risk factor for psychiatric disorders, including schizophrenia [6,7]. Additionally, we identified an exonic deletion in *RELN* in a Japanese patient with schizophrenia through high-resolution copy number variation analysis, which was accompanied by a decreased serum level of reelin [8,9]. Several studies have also reported that reelin protein and mRNA expression levels are reduced by approximately 50% in brain extracts from patients with schizophrenia and autism [10–13]. These findings highlight the importance of reelin in the brain and its strong correlation with psychiatric disorders.

In reelin signaling, reelin directly binds to the extracellular domains of very low-density lipoprotein receptor (VLDLR) and apolipoprotein E receptor 2 (ApoER2) [14,15]. This interaction triggers tyrosine phosphorylation of the cytosolic adaptor protein Disabled homolog 1 (Dab1) by the Src family tyrosine kinases (SFKs), Src and Fyn [16–18], and the kinases themselves undergo autoactivation in the presence of phosphorylated Dab1 [19]. Phosphorylated Dab1 serves as a hub in the reelin signaling cascade, from which several critical downstream pathways diverge, mediating diverse functions of reelin, including neuronal migration [20], dendritic growth and branching [21], actin dynamics [22], dendritic spine development [5], and synaptic plasticity [23]. Although several core elements of the reelin signaling pathway have been identified [4], these elements alone do not fully explain all reelin functions. Therefore, additional downstream molecular candidates that contribute to reelin function remain to be identified.

*Reeler* is a spontaneous autosomal recessive mutation in mice that leads to abnormal brain development [24]. Among the various *reeler* mutations, the “Jackson” strain (*Reln^rl−J^*) exhibits a complete loss of reelin production [25], whereas the “Orleans” strain (*Reln^rl-Orl^*) produces a C-terminal truncated reelin protein that results in defective secretion into the extracellular matrix [26]. Both *Reln^rl−J^*and *Reln^rl-Orl^* mutant mice exhibit abnormal brain layer formation [26,27] along with aberrant behaviors, including heightened anxiety, cognitive dysfunction, impaired social novelty [8,28–30], and defects in spine development and synaptic function [28,31].

To identify molecules acting downstream of the reelin receptor-Dab1 signaling pathway that might contribute to deficits observed in *Reln^rl-Orl^*mice [8], we conducted a comprehensive proteomic analysis to screen for proteins with altered tyrosine phosphorylation levels in the brains of *Reln^rl-Orl^* mice. Twinfilin-1 (Twf1) emerged as a strong candidate from our proteomic analysis owing to its undetectable tyrosine phosphorylation levels in homozygous *Reln^rl-Orl/rl-Orl^* mice. Twf1 is a member of the actin-depolymerizing factor homology (ADF-H) domain superfamily and plays an essential role in regulating actin dynamics by interacting with monomeric actin and capping protein (CP) [32]. Here, we investigated reelin-induced Twf1 phosphorylation and its contribution to synaptic structure and function.

## 2. Results

### 2.1. Twf1 is a downstream effector of reelin signaling

Tyrosine phosphorylation plays a crucial role in reelin signaling [17–19]. Therefore, as a first step in screening for reelin signaling molecules, we conducted a comprehensive proteomics analysis using brain lysates from wild-type (WT) and *reeler* mice. Silver staining after immunoprecipitation with an anti-phosphotyrosine (pY) antibody revealed several proteins with decreased tyrosine phosphorylation levels in brain lysates prepared from homozygous *Reln^rl-Orl/rl-Orl^*and heterozygous *Reln^rl-Orl/+^* mice compared to their WT littermates (Fig. 1A). From the results of the liquid chromatography-tandem mass spectrometry (LC-MS/MS) analysis (Table S1), several potential downstream candidates were identified based on the following criterion: gene dose-dependent changes, with undetectable tyrosine phosphorylation levels in homozygous *Reln^rl-Orl/rl-Orl^*mice and an approximately 50% reduction in heterozygous *Reln^rl-Orl/+^* mice compared to their WT littermates (Table 1). Among these candidates, we focused on Twf1, an actin-depolymerizing factor that plays a crucial role in regulating actin dynamics, as n-cofilin, an actin-polymerizing protein, has already been implicated in reelin signaling [22]. We further validated the LC-MS/MS data through western blot analysis using an anti-Twf1 antibody (Fig. 1A). To determine whether Twf1 can be phosphorylated by SFKs, we co-transfected Myc-Twf1 with GFP-Src or GFP-Fyn into human embryonic kidney 293T (HEK293T) cells. Immunoprecipitation and western blot analysis revealed that both GFP-Src and GFP-Fyn significantly enhanced Twf1 tyrosine phosphorylation, with GFP-Src demonstrating a markedly stronger effect (Fig. 1B). According to the PhosphoSitePlus® database (v6.7.9), tyrosine 309 (Y309) on Twf1 has been identified as a phosphorylation site in high-throughput mass spectrometry studies. To further identify the specific phosphorylation site on Twf1, phospho-resistant Twf1 mutants, Twf1 Y137F and Twf1 Y309F, were co-transfected with GFP-Src into HEK293T cells. Although the Twf1 Y137F mutation significantly reduced Twf1 tyrosine phosphorylation, the Twf1 Y309F mutation completely abolished it (Fig. 1C). These results suggest that Twf1 is a downstream target of reelin signaling and that Src regulates Twf1 phosphorylation at Y309.

**Fig. 1.**
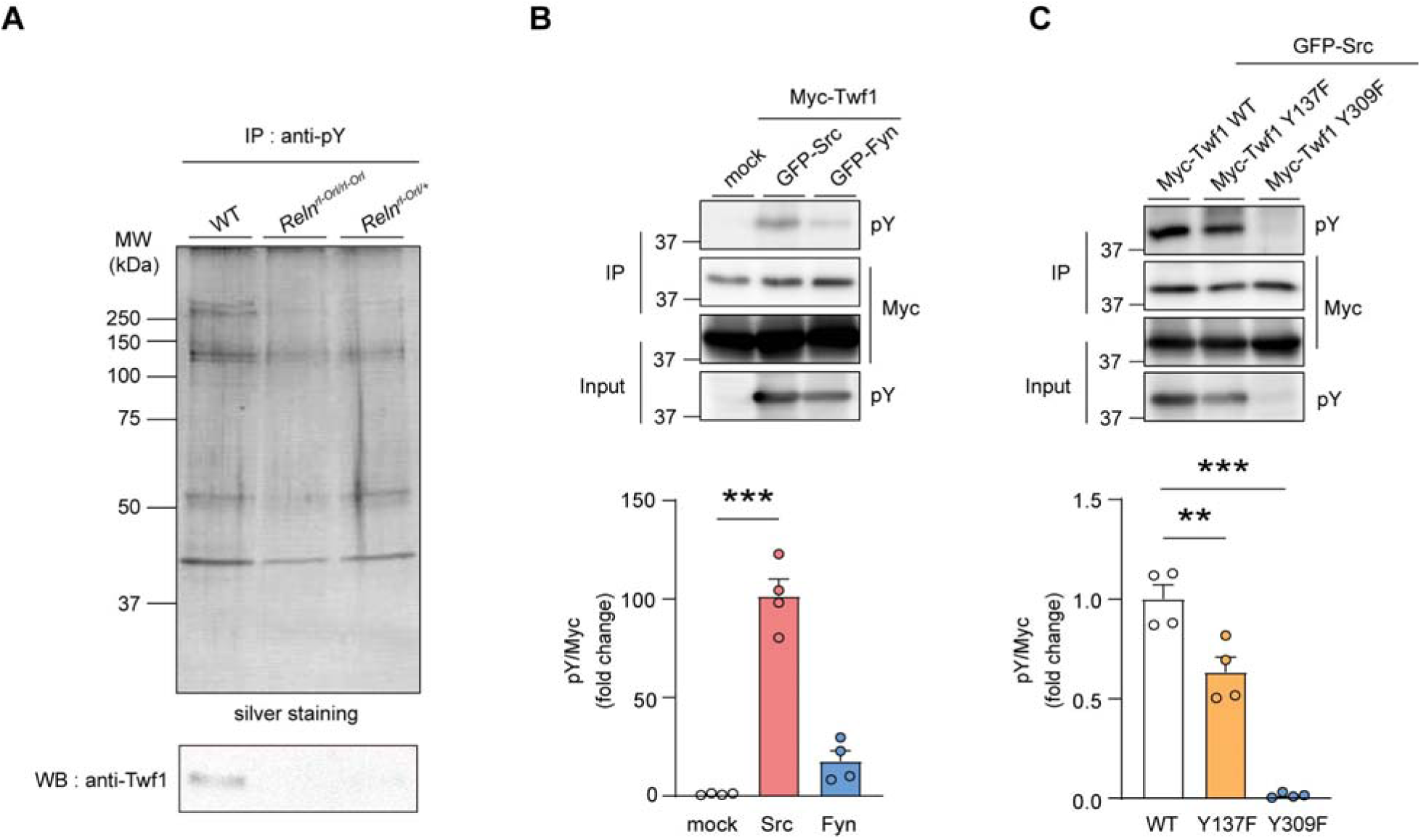
Twf1 is a downstream candidate in the reelin signaling pathway. (A) Upper panel: Silver staining of anti-phosphotyrosine (pY) antibody co-immunoprecipitates from brain lysates of wild-type (WT), homozygous *Reln^rl-Orl/rl-Orl^*, and heterozygous *Reln^rl-Orl/+^* mice. Lower panel: Western blot analysis of anti-pY antibody co-immunoprecipitates using anti-Twf1 antibody. IP, immunoprecipitation; WB, western blotting. (B) Phosphorylation of Twf1 by Src and Fyn kinases. Myc-Twf1 was co-transfected with GFP-Src or GFP-Fyn into human embryonic kidney (HEK 293T) HEK293T cells. Western blot analysis of anti-Myc antibody co-immunoprecipitates using an anti-pY antibody. *** P < 0.001 vs. mock group, Tukey’s multiple comparisons test (n = 4 independent experiments). (C) Identification of Twf1 phosphorylation site by Src. Phospho-resistant mutants Myc-Twf1 Y137F or Myc-Twf1 Y309F were co-transfected with GFP-Src into HEK293T cells. Western blot analysis of anti-Myc antibody co-immunoprecipitates using anti-pY antibody. *** P < 0.001, ** P < 0.01 vs. Myc-Twf1 WT group, Tukey’s multiple comparisons test (n = 4 independent experiments). All data represent the mean ± standard error of the mean (SEM).

**Table 1.** Identification of Twinfilin-1 (Twf1) as a downstream candidate in the reelin signaling pathway via liquid chromatography-tandem mass spectrometry.

| Description | MW<br>(kDa) | Score |  |  | Coverage |  |  |
| --- | --- | --- | --- | --- | --- | --- | --- |
|  |  | WT | *Reln*<sup>rl-Orl/rl-Orl</sup> | *Reln*<sup>rl-Orl/+</sup> | WT | *Reln*<sup>rl-Orl/rl-Orl</sup> | *Reln*<sup>rl-Orl/+</sup> |
| Myosin-14 | 228.4 | 2263.95 | N/D | 1252.35 | 18.6 | N/D | 10.35 |
| **Twinfilin-1** | **40.1** | **1397.76** | **N/D** | **719.90** | **26.29** | **N/D** | **24.57** |
| PH and SEC7 domain-containing protein 3 | 114.7 | 1979.78 | N/D | 1245.61 | 15.81 | N/D | 13.60 |
| Unconventional myosin-VI | 146.3 | 1076.00 | N/D | 713.58 | 14.23 | N/D | 12.73 |
| Tenascin | 231.7 | 1063.41 | N/D | 701.89 | 11.56 | N/D | 9.24 |
| Actin-binding LIM protein 2 | 68.1 | 1304.85 | N/D | 602.60 | 22.39 | N/D | 14.54 |
| Coronin-1B | 53.9 | 1075.07 | N/D | 362.66 | 31.82 | N/D | 24.59 |

MW, molecular weight.

### 2.2. Recombinant reelin induces Twf1 phosphorylation in primary cultured cortical neurons via Src activation

Previous studies have reported that reelin stimulation of primary cultured neurons induces intracellular Dab1 and Src tyrosine phosphorylation [16,19]. To examine whether reelin also stimulates Twf1 phosphorylation, primary cultured cortical neurons from WT mice were treated with recombinant reelin on day *in vitro* (DIV) 14 (DIV14), a stage when cultured neurons undergo significant development and maturation [33]. A reelin mutant in which lysines (K) at positions 2360 and 2467 are substituted with alanine (A) (K2360A/2467A; K2A) was used as a negative control that is incapable of binding reelin receptors and activating downstream signaling [34,35]. Phosphorylated proteins were detected using an anti-pY antibody in immunoprecipitation and analyzed by western blotting. Treatment of primary cultured cortical neurons with WT reelin not only activated Src phosphorylation (Fig. 2A) but also induced Dab1 tyrosine phosphorylation (Fig. 2B), consistent with previous reports [16,18,19]. In contrast, K2A reelin failed to induce phosphorylation of Src and Dab1 (Fig. 2A, B). Notably, WT reelin, but not K2A reelin, also induced Twf1 phosphorylation in primary cultured cortical neurons from WT mice (Fig. 2C). These results were further supported by experiments using cultured cortical neurons from reelin-deficient *reeler* mice (Fig. S1A–C).

**Fig. 2.**
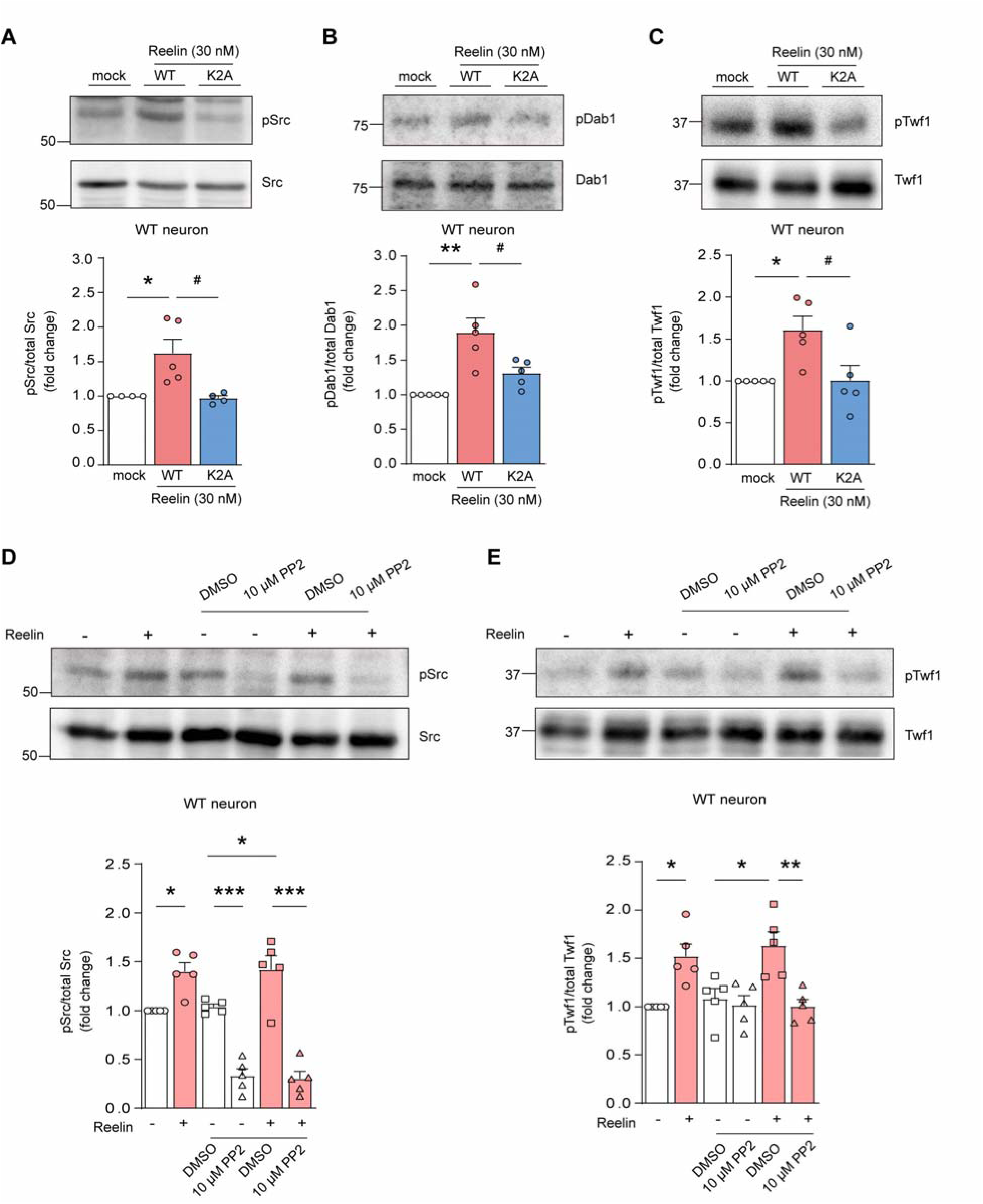
Recombinant reelin induces Twf1 phosphorylation in primary cultured cortical neurons via Src activation. (A–C) Effects of WT and mutant K2A reelin on the phosphorylation levels of (A) Src, (B) Disabled homolog 1 (Dab1), and (C) Twf1 in primary cultured cortical neurons. Cultured neurons at 14 days *in* vitro (DIV14) were treated with WT or K2A reelin at 30 nM for 15 minutes. ** P < 0.01, * P < 0.05 vs. mock group; ^#^ P < 0.05 vs. WT reelin group, Tukey’s multiple comparisons test (n = 4–5 independent experiments). (D and E) Effect of Src inhibitor PP2 on reelin-induced (D) Src and (E) Twf1 phosphorylation. Cultured neurons were pre-treated with PP2 at 10 μM for 1 hour and then stimulated with reelin at 30 nM for 15 minutes. *** P < 0.001, ** P < 0.01, * P < 0.05, Tukey’s multiple comparisons test (n = 5 independent experiments). All data represent the mean ± SEM.

Given that Twf1 phosphorylation was induced by Src overexpression in HEK293T cells (Fig. 1B), we investigated whether reelin-induced Twf1 phosphorylation in primary cultured neurons depends on Src activation. To test this, we used a commercially available tyrosine kinase inhibitor, PP2, which inhibits Src phosphorylation in an ATP-competitive manner [36,37]. Cultured neurons were pre-treated with 10 μM PP2 for 1 h, and the inhibition efficiency of PP2 on reelin-induced Src phosphorylation was validated (Fig. 2D). Western blot analysis revealed that PP2 effectively suppressed reelin-induced Twf1 phosphorylation (Fig. 2E).

Taken together, these results indicate that Twf1 phosphorylation is involved in reelin signaling and depends on reelin binding to its receptors as well as subsequent intracellular Src activation.

### 2.3. Phospho-resistant Twf1 Y309F reduces CP interaction and impairs actin dynamics in HEK293T cells

To elucidate the functional significance of Twf1 phosphorylation, particularly its effect on actin dynamics, we transfected HEK293T cells with one of six plasmid constructs: mock, human TWF1 (hTWF1) WT, hTWF1 WT + Src WT, hTWF1 WT + kinase-dead (KD) Src (Src KD), hTWF1 WT + constitutively active (CA) Src (Src CA), or hTWF1 Y309F + Src CA. hTWF1 was tagged with Flag to facilitate its extraction via immunoprecipitation, followed by western blotting using anti-Flag, anti-β actin, and anti-CP antibodies. First, proteins from transfected cells were separated on a Phos-tag gel, which efficiently distinguishes phosphorylated from non-phosphorylated proteins [38]. Western blot analysis confirmed that hTWF1 WT was strongly phosphorylated in HEK293T cells when co-expressed with Src CA (Fig. 3A). As expected, hTWF1 Y309F remained unphosphorylated even in the presence of Src CA. Next, we evaluated actin and CP binding to hTWF1, as Twf1 regulates actin dynamics through its interaction with actin and CP [39–41]. Notably, the interaction between hTWF1 and CP in the presence of Src CA was markedly reduced when hTWF1 Y309F was expressed instead of hTWF1 WT (Fig. 3B, C), suggesting that Twf1 phosphorylation enhances its interaction with CP. A similar trend was observed for the hTWF1–actin interaction, though the change was not statistically significant owing to high variability (Fig. 3B, D).

**Fig. 3.**
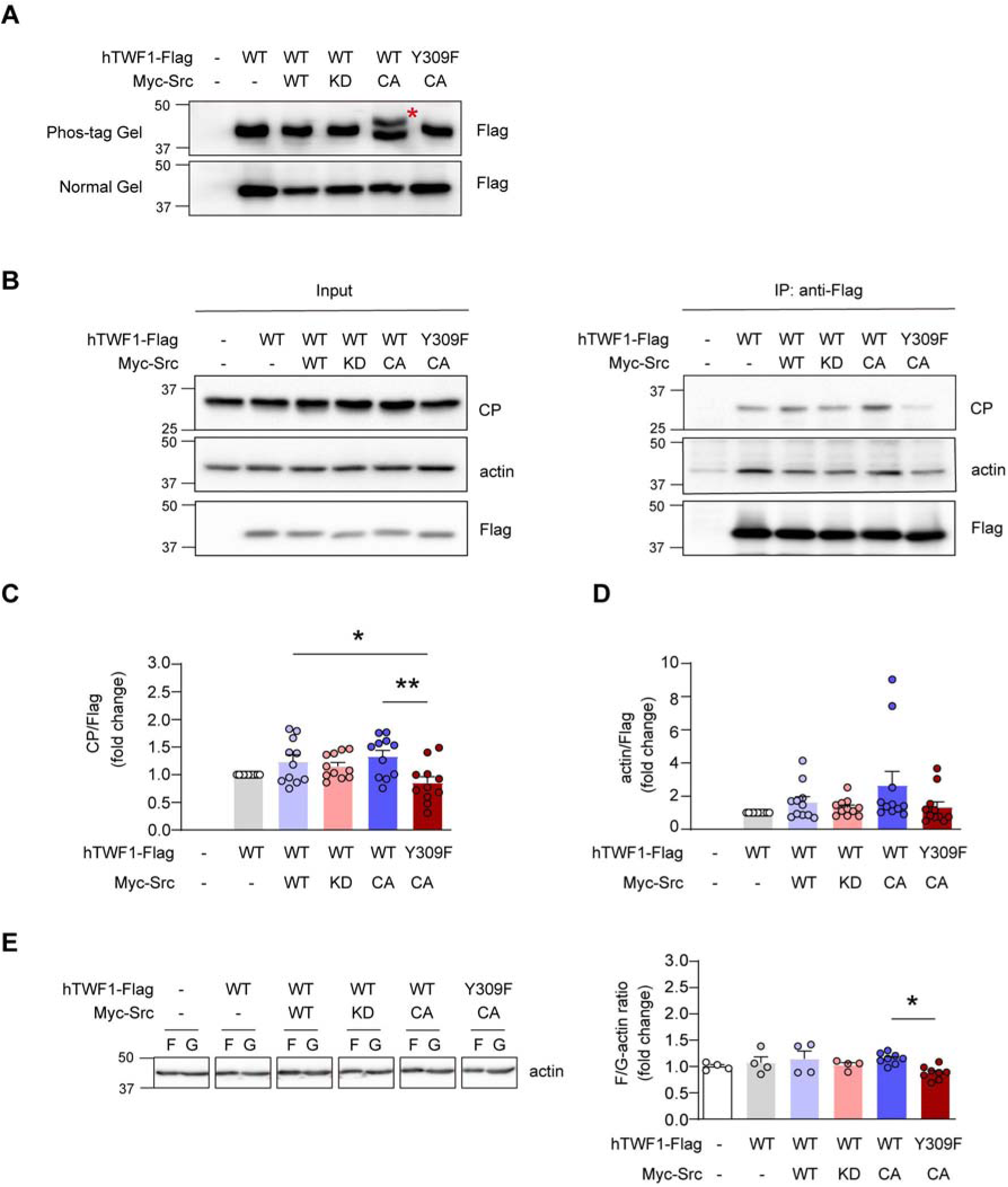
Phospho-resistant Twf1 Y309F reduces capping protein (CP) interaction and impairs actin dynamics in HEK293T cells. (A) Western blot analysis of phosphorylated proteins in HEK293T cells transfected with one of six plasmid constructs: mock, human TWF1 (hTWF1) WT, hTWF1 WT + Src WT, hTWF1 WT + Src KD (kinase-dead), hTWF1 WT + Src CA (constitutively active), or hTWF1 Y309F + Src CA. The red asterisk (*) indicates phosphorylated hTWF1-Flag. (B–D) Assessment of hTWF1 interaction with CP and actin in HEK293T cells transfected with one of the six plasmid constructs. (B) Representative western blot images of anti-Flag antibody co-immunoprecipitates with anti-CP, anti-β actin, and anti-Flag antibodies. Lower panel: Quantification of CP (C) and actin (D) binding to Flag-tagged TWF1. ** P < 0.01, * P < 0.05 vs. hTWF1 Y309F + Src CA group, Tukey’s multiple comparisons test (n = 11 independent experiments). (E) Western blot analysis of F-actin and G-actin in HEK293T cells transfected with the same six plasmid constructs. F, F-actin; G, G-actin. * P < 0.05 vs. hTWF1 Y309F + Src CA group, Tukey’s multiple comparisons test (n = 4 independent experiments). All data represent the mean ± SEM.

Given that Twf1 is an actin-depolymerizing factor that regulates actin dynamics [32], we examined whether Twf1 phosphorylation influences actin polymerization by measuring F-actin and G-actin levels in HEK293T cells transfected with the six plasmid constructs. Consistent with changes in CP– and actin–Twf1 interactions, co-transfection of hTWF1 Y309F with Src CA significantly reduced the F/G-actin ratio compared to co-transfection of hTWF1 WT with Src CA (Fig. 3E). These results suggest that Twf1 phosphorylation at Y309 enhances its interaction with CP, leading to an increase in the F/G-actin ratio.

### 2.4. Expression of phospho-resistant Twf1 Y309F in the medial prefrontal cortex (mPFC) of mice reduces spine head size in neurons

The actin cytoskeleton is highly enriched in dendritic spines, where its dynamic remodeling is essential for spine development and morphological plasticity [42]. Given that the expression of hTWF1 Y309F reduced the F/G-actin ratio in HEK293T cells (Fig. 3E), we examined whether Twf1 phosphorylation influenced dendritic spine structure and morphology. To this end, Twf1 WT, Y309E, Y309F, or an empty EGFP-expressing lentiviral vector (pLLX) was bilaterally microinjected into the mPFC of mice. Dendritic spines were visualized via EGFP immunostaining and analyzed using the Imaris imaging system. Infusion of lentivirus expressing Twf1 WT into the mPFC decreased spine head diameter (Fig. S2A–C), whereas Twf1 Y309F significantly decreased both spine head diameter and volume (Fig. S2A–D). However, when average spine head volume per neuron was analyzed, neurons expressing Twf1 Y309F did not show a statistically significant reduction, although a decreasing trend was observed (Fig. S2E). These data suggest that Twf1 phosphorylation is required for normal spine morphology.

### 2.5. Twf1 phosphorylation regulates its subcellular localization in primary cultured cortical neurons

A previous study reported that Twf1 is enriched in dendritic spines, where its localization influences spine stability [40]. Given that overexpression of the phospho-resistant Twf1 Y309F mutant resulted in reduced neuronal spine size (Fig. S2), we investigated whether Twf1 phosphorylation affects its cellular localization. To test this, we transfected primary cultured cortical neurons with Myc-tagged Twf1 WT, phospho-mimic Twf1 Y309E, or phospho-resistant Twf1 Y309F on DIV18 and analyzed their spine-to-dendritic shaft localization ratios on DIV21. The Myc-fluorescence intensity was normalized to that of co-expressed mCherry [43]. Twf1 Y309E exhibited significantly higher spine enrichment than Twf1 WT and Twf1 Y309F in primary cultured neurons (Fig. 4A, B), suggesting that Twf1 phosphorylation affects its cellular localization by increasing its enrichment in spines.

**Fig. 4.**
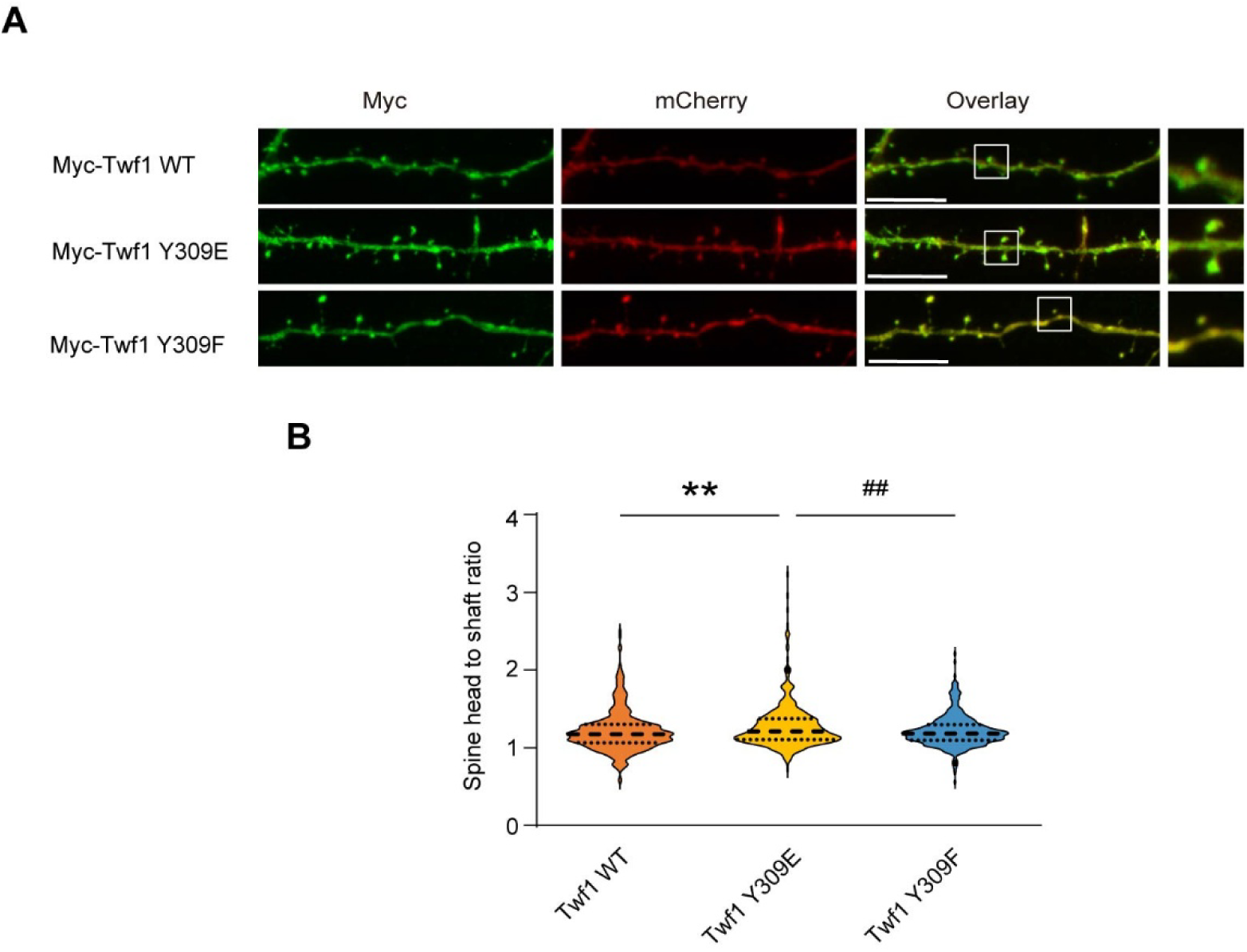
Twf1 phosphorylation regulates its subcellular localization in primary cultured cortical neurons. (A) Representative images of cortical neurons at DIV14 co-expressing mCherry (volume marker) with either Myc-Twf1 WT, phospho-mimic Myc-Twf1 Y309E, or phospho-resistant Myc-Twf1 Y309F. Scale bar: 10 μm. Magnified views of the regions outlined by the white boxes are shown on the far right. (B) Quantification of Twf1 subcellular localization, expressed as the ratio of Myc fluorescence intensity in the spine head to the intensity of an equal-sized area in the adjacent dendritic shaft. Values were normalized to the mCherry intensity in the corresponding regions. n = 375–428 spines from 23–27 dendrites (n = 4 independent experiments). ** P < 0.01 vs. Twf1 WT group; ^##^ P < 0.01 vs. Twf1 Y309E group, Tukey’s multiple comparisons test. All data represent the mean ± SEM.

### 2.6. Homozygous Twf1^Y309F/Y309F^ mice exhibit impaired cognitive function in the visual discrimination (VD) and reversal learning (RL) tasks

To investigate the functional significance of Twf1 phosphorylation *in vivo*, we generated genetically modified mice in which Twf1 phosphorylation at Y309 was disrupted (*Twf1^Y309F^* mice), using a CRISPR/Cas9 knock-in strategy (Fig. S3A–C). These mice were developed on a C57BL/6J genetic background, a standard murine strain for general behavioral analyses.

The body weight of *Twf1^Y309F^* mice was comparable to that of their WT littermates (Fig. S3D), suggesting that decreased Twf1 phosphorylation levels did not affect general growth. We next conducted global behavioral analyses using the following four behavioral tests: locomotor, open field, Y-maze, and social interaction tests. No significant differences were observed between WT and *Twf1^Y309F^* mice in these tests (Fig. S4A–H), suggesting that reduced Twf1 phosphorylation does not affect general behavioral functions.

Our previous study reported that abnormal spine formation in the mPFC is associated with cognitive impairment in the VD task [44]. Additionally, we found that mice with *RELN* deletion (*Reln*-del mice) exhibited cognitive deficits in the VD task [45]. To determine whether Twf1 phosphorylation influences spine formation and contributes to the cognitive impairments observed in *Reln*-del mice, we evaluated the cognitive function of *Twf1^Y309F^* mice in the VD task (Fig. 5A).

**Fig. 5.**
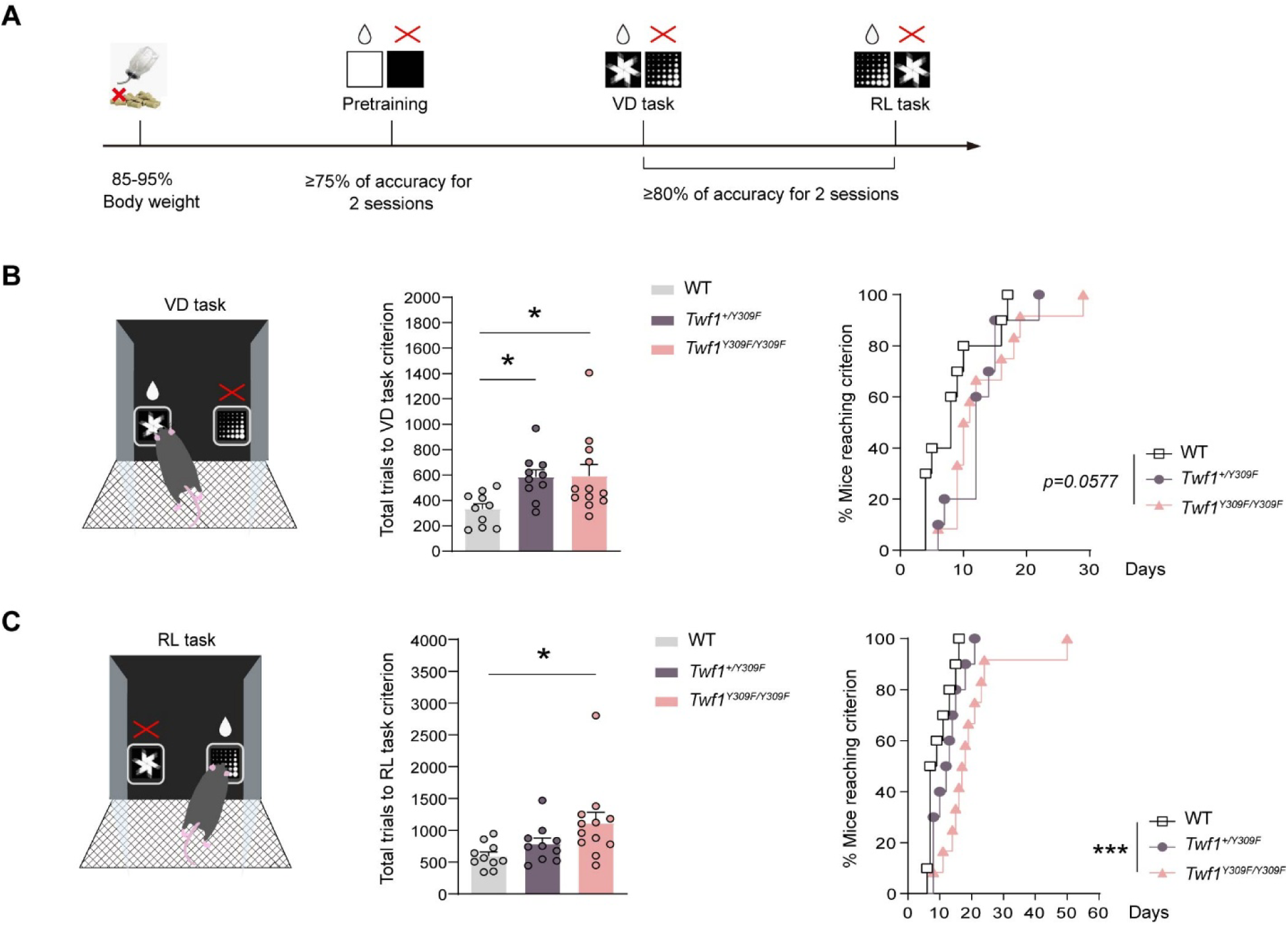
Homozygous *Twf1^Y309F/Y309F^* mice exhibit impaired cognitive function in the visual discrimination (VD) and reversal learning (RL) tasks. (A) Experimental timeline for touchscreen-based behavioral tests. (B) Performance of WT, *Twf1^+/Y309F^*, and *Twf1^Y309F/Y309F^*mice in the VD task. Left: Total number of trials required to reach the VD task criterion. Right: Percentage of mice that reached the VD task criterion over consecutive training days. (C) Performance of WT, *Twf1^+/Y309F^*, and *Twf1^Y309F/Y309F^* mice in the RL task. Left: Total number of trials required to reach the RL task criterion. Right: Percentage of mice that reached the RL criterion over daily training sessions. Group sizes are as follows: WT mice (n = 10, male = 5, female = 5); *Twf1^+/Y309F^* mice (n = 10, male = 5, female = 5); and *Twf1^Y309F/Y309F^*mice (n = 12, male = 6, female = 6). *** P < 0.001, * P < 0.05 vs. WT group, Dunnett’s multiple comparisons test (left) and log-rank test (right). All data represent the mean ± SEM.

The VD task assesses goal-directed discriminative learning and memory [46], while the RL task evaluates choice shifting and relearning ability [47]. Heterozygous *Twf1^+/Y309F^* mice, homozygous *Twf1^Y309F/Y309F^*mice, and their WT littermates showed no significant differences in their ability to complete the pretraining stages (Fig. S4I), suggesting normal visuospatial and motor functions. In the VD task, all mice were required to touch one of two visual stimuli (marble or fan) to receive a reward (Fig. 5B). Homozygous *Twf1^Y309F/Y309F^* mice required significantly more trials to reach the criterion (≥80% correct response for two consecutive days) than WT mice, with a similar trend in heterozygous *Twf1^+/Y309F^* mice (Fig. 5B). Additionally, homozygous *Twf1^Y309F/Y309F^* mice required more training sessions to reach the criterion than WT mice (Fig. 5B). After completing the VD task, mice underwent the RL task, in which reward contingencies were reversed (Fig. 5C). In this task, homozygous *Twf1^Y309F/Y309F^*mice required approximately twice as many training trials as WT mice to reach the criterion (≥80% correct response for two consecutive days) (Fig. 5C). The percentage of mice reaching the criterion was significantly lower in homozygous *Twf1^Y309F/Y309F^*mice, indicating impaired choice-shifting and relearning, likely owing to the increased number of training sessions required to complete the RL task (Fig. 5C). Taken together, these findings indicate that reward-directed discriminative learning and relearning ability were significantly impaired in homozygous *Twf1^Y309F/Y309F^* mice.

### 2.7. Homozygous Twf1^Y309F/Y309F^ mice show abnormal spine morphology and reduced F/G-actin ratio despite normal cortical layer structure

Given that reelin plays a crucial role in brain layer formation and dendritic spine development [4], we investigated whether Twf1 phosphorylation is also involved in these processes by assessing histological changes in the brains of *Twf1^Y309F^* mice. First, we measured the thickness of the forebrain cortex layers using two layer-specific markers: Cux1 for layers II–III and Ctip2 for layers V–VI [48,49]. No significant differences were observed in brain layer thickness among WT, heterozygous *Twf1^+/Y309F^*, and homozygous *Twf1^Y309F/Y309F^* mice (Fig. 6A, B). Therefore, *Twf1^Y309F^* mice did not exhibit apparent abnormalities in cortical layer formation, unlike the *reeler* mutant mice [4,27,50].

**Fig. 6.**
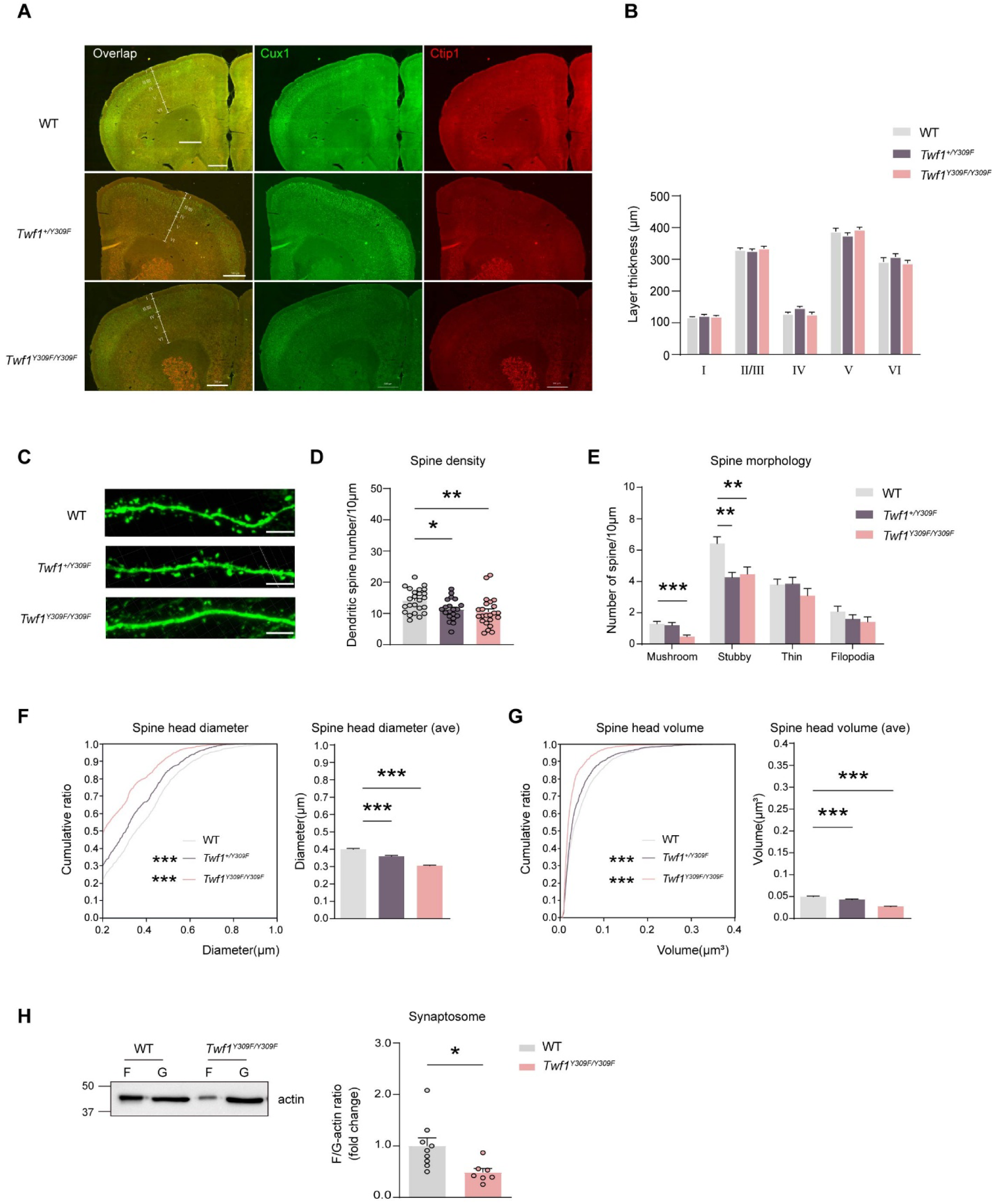
Homozygous *Twf1^Y309F/Y309F^* mice exhibit abnormal spine morphology and reduced F/G-actin ratio despite normal cortical layer structure. (A) Representative images of the cerebral cortex from WT, heterozygous *Twf1^+/Y309F^*, and homozygous *Twf1^Y309F/Y309F^* mice brain sections stained with anti-Cux1 (green) and anti-Ctip2 (red) antibodies. All sections were obtained from 8–10-week-old mice. Scale bar: 500 μm. (B) Quantification of the thickness of layer I, layer II/III (Cux1-dense region), layer IV, layer V (Ctip2-dense region), and layer VI (Ctip2-sparse region). n = 6 mice/group (3–4 slices/mice). (C) Representative images of dendritic spines in the medial prefrontal cortex (mPFC) of WT, *Twf1^+/Y309F^*, and *Twf1^Y309F/Y309F^* mice. Scale bar: 5 µm. (D) Graphical representation of spine density of apical dendrites in mPFC of WT, *Twf1^+/Y309F^*, and *Twf1^Y309F/Y309F^* mice. ** P < 0.01, * P < 0.05 vs. WT group, Dunnett’s multiple comparisons test. e, Graphical representation of the number of mushroom, stubby, thin, and filopodia spines per 10 μm apical dendrites in the mPFC of WT, *Twf1^+/Y309F^*, and *Twf1^Y309F/Y309F^* mice. *** P < 0.001, ** P < 0.01 vs. WT group, Dunnett’s multiple comparisons test. (F) Cumulative ratio (left) and average (right) of spine head diameter in the mPFC of WT, *Twf1^+/Y309F^*, and *Twf1^Y309F/Y309F^* mice (n = 972– 1523 spines from 5 mice). Cumulative ratio, *** P < 0.001, vs. WT group, Kolmogorov– Smirnov test. Average diameter and volume, *** P < 0.001 vs. WT group, Dunnett’s multiple comparisons test. (G) Cumulative ratio (left) and average (right) of spine head volume in the mPFC of WT, *Twf1^+/Y309F^*, and *Twf1^Y309F/Y309F^* mice (n = 972– 1523 spines from 5 mice). Cumulative ratio, *** P < 0.001 vs. WT group, Kolmogorov– Smirnov test. Average diameter and volume, *** P < 0.001 vs. WT group, Dunnett’s multiple comparisons test. (H) F/G-actin ratio in synaptosome fractions from cortical neurons at DIV15 in WT and *Twf1^Y309F/Y309F^* mice. F, F-actin; G, G-actin. * P < 0.05 vs. WT group, WT group: n = 9 wells, *Twf1^Y309F/Y309F^* group: n = 7 wells (3 independent experiments), unpaired Student’s *t*-test. All data represent the mean ± SEM.

Next, we analyzed dendritic spine morphology in pyramidal neurons of the mPFC, a region implicated in reward-directed discriminative learning and relearning processes [47], and in various neurological disorders [51]. To visualize spine morphology, we bilaterally injected an EGFP-expressing lentiviral vector (pLLX) into the mPFC of 8-week-old mice, enabling clear visualization of spine morphology and *in vivo* labeling of layer II/III neurons. The number of dendritic spines in the mPFC was significantly lower in homozygous *Twf1^Y309F/Y309F^*mice than in WT mice (Fig. 6C, D). We classified spine morphology into four subtypes: mushroom, stubby, thin, and filopodia. Mushroom and stubby spines were significantly decreased in homozygous *Twf1^Y309F/Y309F^* mice, while heterozygous *Twf1^+/Y309F^* mice also exhibited a decrease in stubby spines (Fig. 6E). Furthermore, both heterozygous and homozygous *Twf1^Y309F/Y309F^* mice had significantly smaller spine head diameters and volumes (Fig. 6F, G). To examine the biochemical basis of these morphological changes, we measured F-actin and G-actin levels in synaptosomes from cultured neurons derived from WT and homozygous *Twf1^Y309F/Y309F^* mice. As shown in Fig. 6H, the F/G-actin ratio was significantly decreased in synaptosomes from homozygous *Twf1^Y309F/Y309F^*neurons compared to WT neurons, which is consistent with the finding that expression of hTWF1 Y309F in HEK293T cells also reduced the F/G-actin ratio (Fig. 3E).

Lastly, we investigated total Twf1 levels and Twf1-associated proteins, including cofilin and CP, which cooperate with Twf1 to regulate actin polymerization and depolymerization [40,52]. WT, heterozygous, and homozygous *Twf1^Y309F/Y309F^* mice had comparable levels of Twf1, cofilin, and CP (Fig. S5A–D) in the brain, indicating that reduced Twf1 phosphorylation does not affect the overall levels of Twf1 or its associated proteins. Collectively, these findings suggest that Twf1 phosphorylation is critical for regulating the F/G-actin ratio, influencing the number, size, and shape of dendritic spines, but has minimal effects on cortical layer formation *in vivo*.

### 2.8. RELN-del induced pluripotent stem cells (iPSCs) from a patient with schizophrenia exhibit reduced Twf1 phosphorylation and disrupted F-actin dynamics

A previous study has reported that *RELN*-del iPSCs derived from a patient with schizophrenia exhibit impaired reelin signaling and migration defects compared to iPSCs derived from healthy controls [53]. To determine whether *RELN*-del neurons also exhibit impaired Twf1 phosphorylation, we performed immunoprecipitation using an anti-pY antibody, followed by western blotting with an anti-Twf1 antibody. The levels of phosphorylated Twf1 (pTwf1) were decreased in *RELN*-del neurospheres compared to control lines (Fig. S6A). Given our findings that Twf1 phosphorylation is involved in actin dynamics, we next examined the formation of filopodia in the neurospheres, a process that depends on F-actin remodeling [54]. Control neurospheres displayed typical filopodia development, whereas *RELN*-del neurospheres exhibited fewer and shorter filopodia (Fig. S6B–E). These findings confirm that this rare *RELN*-del variant decreases Twf1 phosphorylation and disrupts F-actin dynamics in human neuronal cells.

## 3. Discussion

Reelin exerts multiple critical functions in the developing and adult brain, including regulating neuronal migration [20], dendritic growth [21,55], spine development [5], synapse formation [23,56], and synaptic plasticity [23,57]. Consequently, dysregulation of the reelin signaling pathway has been implicated in various neuropsychiatric disorders, including schizophrenia [4,6,9–13]. Key components of this pathway, such as the reelin receptors VLDLR and ApoER2, the Src family kinases Src and Fyn, and the intracellular adaptor protein Dab1, contribute to most, but not all, reelin-mediated functions [4]. Our study provides precise mechanistic insights into the reelin signaling pathway, identifying Twf1 phosphorylation as a critical downstream component. Tyrosine phosphorylation of Twf1 was significantly reduced in the brains of reelin-deficient *reeler* mutant mice, while stimulation with recombinant reelin strongly induced Twf1 phosphorylation in both WT and *reeler* cortical neurons. Furthermore, by inhibiting Src activation in cortical neurons using PP2, we demonstrated that Src activation is essential for Twf1 phosphorylation.

Twf1 is broadly expressed in mammalian tissues [58], where it regulates actin dynamics through direct interactions with actin monomers, filaments, and CP [59,60], contributing to processes such as cell motility and synaptic endocytosis [61]. It promotes actin filament barbed-end depolymerization and accelerates the uncapping of CP from barbed ends, maintaining actin cytoskeleton treadmilling within the cytoplasm [41,62]. However, until now, little has been known about the functional regulation of Twf1 by tyrosine phosphorylation. We found that reelin activated Src, which in turn regulated Twf1 phosphorylation at Y309 within the C-terminal ADF domain. Furthermore, the phospho-resistant Twf1 mutant, Twf1 Y309F, exhibited reduced interaction with CP and subsequently decreased the F/G-actin ratio in HEK293T cells. A recent study demonstrated that Twf1 functions as a depolymerase at free barbed ends but also promotes filament assembly by facilitating CP dissociation, enabling the resumption of formin-based filament elongation when both CP and formin are present [39]. Based on this model, we speculate that inhibiting Twf1 phosphorylation may disrupt its CP uncapping activity, resulting in a stable association between CP and cellular F-actin arrays. This, in turn, could prevent actin filament elongation or depolymerization, thereby reducing F-actin turnover rates. The different phosphorylation states of Twf1 may enable it to participate in distinct regulatory processes that govern actin cytoskeleton treadmilling. However, further studies are required to confirm this hypothesis.

Dendritic spines are small actin-rich protrusions extending from the dendritic shaft, primarily composed of the actin cytoskeleton [63]. Dynamic actin remodeling is essential for dendritic spine development and morphological plasticity [42]. Actin-binding proteins, such as cofilin and Twf1, regulate this process by modulating actin dynamics [64,65]. Twf1 is highly expressed in the mammalian brain [58,59], where it cooperates with CP to remodel the actin cytoskeleton in dendritic spines [40]. Knockdown of Twf1 in cultured neurons leads to decreased spine density and increased filopodia density and instability [40]. Our study highlights the critical role of Twf1 phosphorylation at Y309 in spine development and morphology. We demonstrated that expression of the phospho-resistant Twf1 Y309F mutant in the brain resulted in smaller spine heads, potentially as a result of decreased actin dynamics. This finding phenocopies Twf1 knockdown in spines, resembling the effects of CP knockdown [40,66], and aligns with previous reports identifying Twf1 as an uncapping ligand of CP that promotes actin turnover [41]. Moreover, the phospho-mimic Twf1 Y309E mutant was enriched in dendritic spines, where CP is also concentrated [40]. This suggests that interaction with CP serves as a key mechanism for dynamically regulating pTwf1 activity and its association with actin barbed ends in spines. Further studies are required to determine whether Twf1 phosphorylation regulates actin dynamics in non-neuronal cells.

Several *Reln* mutant mouse models have been developed to investigate reelin signaling in the brain, from development to postnatal stages [8,45,50]. To elucidate the pathophysiological role of reelin–Twf1 signaling *in vivo*, we generated a *Twf1^Y309F^* mouse model to mimic the decreased Twf1 phosphorylation observed in *reeler* mutants and compared our findings with previous *reeler-*associated models.

*Reln^rl-Orl/^*^+^ mice exhibited anxiety-like behavior and social dysfunction [8], while *Reln^del/+^* mice showed abnormalities in social novelty and cognitive function [45,50], both linked to impairments in reelin signaling. Through comprehensive behavioral analyses, we identified cognitive dysfunction in *Twf1^Y309F^* mice, suggesting that they partially mimic behavioral deficits observed in *reeler*-associated models. Touchscreen-based behavioral tasks have been developed as an effective translational approach for assessing higher cognitive functions associated with neuropsychiatric disorders in rodents, enabling cross-species comparisons and evaluations of pharmacological interventions [67]. These computerized operant behavioral tests offer high reproducibility, low variability, and increased sensitivity to cognitive abnormalities in mice [67]. Clinically, impaired cognitive function is a hallmark of patients with schizophrenia with a *RELN* exonic deletion [8], consistent with our findings in *Twf1^Y309F^*mice.

Reelin deficiency disrupts brain layer formation, a characteristic of the *reeler* phenotype [4], leading to neuronal ectopia in laminated structures, such as the cerebral and cerebellar cortices [50,68]. To investigate whether the reelin–Twf1 signaling influences brain layer formation, we examined *Twf1^Y309F^* mouse brain histology. Notably, these mice exhibited normal brain layer formation and structure, suggesting that reelin– Twf1 signaling is not essential for neuronal migration during development. Most heterozygous reelin mutant mice exhibited behavioral deficits without clear histological abnormalities [8,28,30,69,70]. These deficits may stem from subtle changes in neuronal plasticity and spine formation rather than gross structural defects. Additionally, dendritic growth is disrupted in both homozygous and heterozygous *reeler* mice, despite normal neuronal positioning in heterozygous mutants [55,71–73]. These findings suggest that dendritic abnormalities are not merely a secondary consequence of neuronal mispositioning, underscoring the need for further investigation into reelin–Twf1 signaling in spine development and function.

Reelin signaling plas a crucial role in spine formation and synaptic plasticity [23,74]. Several core components of the reelin pathway, including VLDLR, ApoER2, SFKs, and Dab1, regulate dendritic spine growth and maturation [71]. Reelin also mediates tyrosine phosphorylation and enhances N-methyl-D-aspartate receptor function in neurons in a Dab1- and SFK-dependent manner, promoting spine enlargement [56,57,75]. In *reeler* mice, decreased spine density and abnormal spine morphology have been reported [8,31,76]. Similarly, *Twf1^Y309F^*mice exhibited a significant decrease in spine density and abnormal morphology in cortical pyramidal neurons. These findings suggest that deficits in reelin–Twf1 signaling impair spine development and maturation, potentially contributing to deficits in long-term potentiation and cognitive dysfunction.

Studies on human brains from patients with neuropsychiatric and neurodegenerative disorders highlight the importance of reelin signaling in disease etiology [4]. It is crucial to examine whether the reelin–Twf1 pathway also plays a role in human brain disorders such as lissencephaly [77], autism [78], schizophrenia [6,9–13,79,80], bipolar disorder [11], mood disorder [12], and Alzheimer’s disease [81], all of which have been linked to reelin dysfunction. Our previous trajectory analysis of highly homologous neurons carrying a rare *RELN* variant, including neurospheres from the patient-derived iPSCs, provides a valuable approach to understanding the mechanisms underlying microstructural defects in the brains of individuals with schizophrenia [53,82]. Notably, Twf1 phosphorylation was markedly decreased in neurospheres derived from iPSCs of a Japanese patient with schizophrenia carrying a *RELN*-del variant, and these cells exhibited a morphological defect resembling the spine abnormalities observed in *Twf1^Y309F^* mutant mice. Further studies are needed to establish a definitive link between the reelin–Twf1 signaling pathway and the etiology of other neuropsychiatric disorders.

Our study has some limitations. First, although Src is implicated in Twf1 phosphorylation, it remains unclear whether Src directly phosphorylates Twf1. Immunostaining for pTwf1 would provide stronger evidence for its localization in neurons and enable direct visualization of reelin-induced Twf1 phosphorylation during spine development and morphological changes. Additionally, while Twf1 has been shown to accelerate CP dissociation [39], the precise mechanism remains unclear due to the lack of simultaneous visualization of both proteins at barbed ends. Future studies integrating biochemical, cellular, and neuroscience approaches are necessary to fully elucidate the reelin–Twf1 signaling pathway, its role in synapse development and plasticity, and its potential contributions to synaptic dysfunction and neurological disorders.

In conclusion, our study identified a pathway in which Twf1 functions as a downstream effector of reelin signaling to regulate actin dynamics and spine development. Our findings suggest that targeting reelin signaling or Twf1 phosphorylation may offer promising therapeutic strategies for neurodevelopmental and neuropsychiatric disorders associated with synaptic dysfunction.

## 4. Materials and methods

### 4.1. Animals

C57BL/6J mice were obtained from Japan SLC Inc. (Hamamatsu, Japan). *Reln^rl-Orl^* mice (RBRC00063) were provided by the RIKEN BioResource Center (Tsukuba, Japan) and maintained in our laboratory. *Reln^rl−J^*mice (B6C3Fe-a/a-rl) were purchased from The Jackson Laboratory (Bar Harbor, ME, USA). *Twf1^Y309F^* mice were generated via CRISPR/Cas9 as previously described [83–86]. Guide RNAs were designed to target exon 9 (NM_008971.5) of the mouse *Twf1* gene. WT, heterozygous (*Twf1^+/Y309F^*), and homozygous (*Twf1^Y309F/Y309F^*) mice were obtained by intercrossing heterozygous *Twf1^+/Y309F^* males and females, with WT littermates serving as controls. All procedures adhered to the guidelines established by the Institutional Animal Care and Use Committee of Nagoya University, the *Guiding Principles for the Care and Use of Laboratory Animals* set forth by the Japanese Pharmacological Society, and the *National Institutes of Health Guide for the Care and Use of Laboratory Animals*.

### 4.2. Generation of Twf1^Y309F^ mice

#### 4.2.1. Chemical synthesis of CRISPR RNA (crRNA), trans-activating RNA (tracrRNA), and donor DNA

*Twf1-*crRNA (5′-CAAUGGGGAUGAGCUGACUGguuuuagagcuaugcuguuuug-3′), tracrRNA (5′-AAACAGCAUAGCAAGUUAAAAUAAGGCUAGUCCGUUAUCAACUUGAAAAAGUGGCACCGAGUCGGUGCU-3′), and donor DNA (5′-TGAAGGTTCCCTGACGTTGTTTAACCCCGGTGGTGCCTTCCTCTCTTGCAGATTGA GATAGACAATGGGGATGAGCTGACTGCAGATTTCCTGTTCGATGAAGTCCACCCCAAGCAGCATGCACATAAGCAGAGT-3′) were chemically synthesized and purified via PAGE (Fasmac, Atsugi, Japan).

#### 4.2.2. Pronuclear injection

Injection mixtures were prepared following a previously established method [83]. To generate *Twf1^Y309F^* mice via protein injection, a mixture containing Cas9 protein, *Twf1-* crRNA, tracrRNA, and donor single-stranded DNA (ssDNA) was prepared in Tris-EDTA buffer at the final concentrations of 100 ng/μL, 0.61 pmol/μL, 0.61 pmol/μL, and 10 ng/μL, respectively. Cas9 protein was obtained from New England Biolabs (Ipswich, MA, USA; M0386S). The donor ssDNA, encoding a FLAG tag and a stop codon and targeting exon 9 of the mouse *Twf1* gene (NM_008971.5), was chemically synthesized by Fasmac Inc. (Japan). The mixture was incubated at 37 °C for at least 15 minutes before microinjection into the pronuclei of one-cell-stage zygotes, which were obtained from C57BL/6J mice (Charles River Laboratories, Wilmington, MA, USA).

#### 4.2.3. PCR screening and genotyping

To screen *Twf1^Y309F^* embryos generated by CRISPR-mediated editing, genomic DNA was extracted from tail samples using proteinase K digestion, followed by standard phenol extraction. PCR amplification was performed using KOD FX Neo polymerase (TOYOBO, Osaka, Japan), and the resulting products were analyzed via electrophoresis on a 2% agarose gel. The primer sequences used for *Twf1^Y309F^*detection were 5′-GGGCCATCGCTGGCCTAGCCCTGTTGTAG-3′ and 5′-GAGCATGCTCAAAGCAATCAGCTGCCCG-3′. PCR products were further cloned using the Zero Blunt TOPO PCR Cloning Kit (Life Technologies) and subjected to sequencing analysis as described previously [83]. After establishing the *Twf1^Y309F^*mouse strain, PstI restriction enzyme digestion was used to distinguish WT and knock-in alleles. Digestion of the PCR product from the knock-in allele resulted in cleavage, producing a shorter DNA fragment detectable by electrophoresis, whereas the WT allele remained uncut.

### 4.3. Human participants

The iPSCs used in this study were derived from three individuals: two healthy Japanese donors, a 30-year-old man (CON1) and a 65-year-old woman (CON2), and one 58-year-old male patient with schizophrenia (RELN) [53,82]. All subjects provided written informed consent. The ages listed for the participants correspond to the time of blood sampling for iPSC generation. Generation and application of human iPSCs were approved by the Nagoya University Ethics Committee (approval number: 2012–0184).

### 4.4. Sample preparation

#### 4.4.1. Tissue samples

Mouse brain sections were lysed in ice-cold lysis buffer (25 mM Tris-HCl [pH 7.4], 150 mM NaCl, 1 mM EDTA, 1% NP-40, 0.1% sodium deoxycholate, 0.1% SDS) supplemented with a complete protease inhibitor cocktail (Roche, Basel, Switzerland) and phosSTOP phosphatase inhibitors (Roche). The lysates were sonicated on ice and centrifuged at 15,000 × *g* for 10 minutes at 4 °C. Protein concentrations were measured using a DC Protein Assay Kit (Bio-Rad Laboratories, Hercules, CA, USA) and adjusted to uniform concentrations in lysis buffer.

#### 4.4.2. Cell culture samples

Cultured cells were collected and lysed in the same lysis buffer for 30 minutes on ice with occasional mixing. The lysates were clarified via centrifugation at 15,000 × *g* for 10 minutes at 4 °C. Protein concentrations were measured using the DC Protein Assay Kit before adjustment to uniform concentrations in lysis buffer.

### 4.5. Immunoprecipitation

Immunoprecipitation was performed using Protein G Sepharose 4 Fast Flow (Cytiva, Marlborough, MA, USA) following the manufacturer’s instructions with minor modifications. Briefly, equal amounts of lysate (500–1000 µg) were incubated with 2–4 µg antibody for 2–4 hours at 4 °C with gentle rotation. Immune complexes were captured by adding 30 µL of agarose beads, followed by an additional 1-hour incubation at 4 °C. The beads were washed three times with ice-cold lysis buffer to minimize nonspecific binding. For LC-MS/MS analysis, immunoprecipitated proteins were eluted with 1M NaCl elution buffer and processed for in-solution digestion. For western blotting, the co-immunoprecipitates were eluted by boiling the beads in 1× sodium dodecyl sulfate (SDS) sample buffer for 5 minutes at 95 °C. The proteins were separated via SDS-polyacrylamide gel electrophoresis (SDS-PAGE) and analyzed via western blotting. Input samples (10% of lysate) were used as loading controls.

### 4.6. Silver staining

Immunoprecipitated proteins were subjected to SDS-PAGE and stained using the SilverQuest Silver Staining Kit (Thermo Fisher Scientific, San Jose, CA, USA) according to the manufacturer’s instructions. Briefly, after electrophoresis, the gel was fixed, microwaved at 700 W for 30 seconds, and agitated for 5 minutes. It was then washed with 30% ethanol, microwaved, and agitated again. The gel was incubated in a sensitizing solution, microwaved, and shaken before being washed twice with ultrapure water. For staining, the gel was incubated in staining solution, microwaved, and gently agitated. The staining solution was discarded, and the gel was briefly washed before incubation in developing solution for 5 minutes. Once the bands reached the desired intensity, stop solution was added, and agitation continued for 10 minutes until the color changed from pink to clear. The gel was finally washed with ultrapure water for 10 minutes before analysis.

### 4.7. In-solution digestion and LC-MS/MS analysis

Protein interactions were identified as previously described [87]. Briefly, the eluted proteins were reduced with 5 mM dithiothreitol for 30 minutes and alkylated with 10 mM iodoacetamide for 1 hour in the dark. The proteins were digested with Trypsin/Lys-C (Promega, Madison, WI, USA) at 37 °C for 16 hours. Nanoelectrospray LC-MS/MS was performed using an LTQ Orbitrap mass spectrometer (Thermo Fisher Scientific, Waltham, MA, USA). Peptide and protein identification was performed via Mascot (Matrix Science Inc., Boston, Massachusetts, USA) and X!Tandem (The GPM), searching against the Swiss-Prot and NCBI databases. Mascot parameters were set as follows: variable modifications—carbamidomethylation of cysteine and oxidation of methionine; mass values: monoisotopic; protein mass: unrestricted; precursor mass tolerance: ±10 ppm; fragment mass tolerance: ±0.8 Da; max missed cleavages: 1; and instrument type: ESI-TRAP. Protein validation was performed using Scaffold (Proteome Software, Portland, OR, USA), with candidate proteins required to contain at least two identified peptides and a probability greater than 95%.

### 4.8. Western blotting

Western blotting was performed as described previously with minor modifications [44,88]. Protein samples (20–40 µg) were separated on 10% SDS-polyacrylamide gels and transferred onto polyvinylidene difluoride membranes (Millipore, Burlington, MA, USA). After transfer, the membranes were blocked with Blocking One-P (Nacalai Tesque, Inc., Kyoto, Japan) for 1 hour at room temperature, followed by overnight incubation at 4 °C with primary antibodies diluted in Can Get Signal Solution 1 (Toyobo, Osaka, Japan). The next day, membranes were washed three times with Tris-buffered saline containing 0.05% Tween 20 and incubated for 1 hour at room temperature with horseradish peroxidase-conjugated secondary antibodies diluted in Can Get Signal Solution 2 (Toyobo, Osaka, Japan). Protein bands were detected using ECL Prime Western Blotting Detection Reagents (GE Healthcare, Chicago, IL, USA) and analyzed with a LuminoGraph I imaging system (Atto Instruments, Tokyo, Japan). Western blotting data were expressed as relative fold changes in protein expression, normalized to control levels. Antibodies used for western blotting are listed in Table S2.

### 4.9. Quantitative real-time reverse transcription PCR (qRT-PCR)

qRT-PCR was performed using the 7300 Real-Time PCR System (Applied Biosystems, Foster City, CA, USA). Each 25 μL reaction contained 12.5 μL of Power SYBR Green PCR Master Mix (Applied Biosystems), 0.5–1.0 µg of cDNA, and 0.5 μM of primers for each target gene. All reactions were performed in duplicate, including no-template controls and RT samples. The PCR cycling conditions were as follows: 50 °C for 2 minutes, 95 °C for 10 minutes, followed by 60 cycles of 95 °C for 15 seconds and 60 °C for 1 minute. Primer sequences used in the qRT-PCR are listed in Table S3. Data were analyzed using the 2^−ΔΔCT^ method [89].

### 4.10. Cell culture and transfection

HEK293T cells (ATCC, Manassas, VA, USA) were cultured in Dulbecco’s modified Eagle medium (DMEM; Sigma-Aldrich, Burlington, MA, USA) supplemented with 10% fetal bovine serum (Gibco, Waltham, MA, USA), 100 units/mL of penicillin, 0.1 mg/mL of streptomycin, and 0.25 μg/mL of amphotericin B (Nacalai Tesque, Inc.). Cells were maintained at 37 °C under humidified air with 5% CO_2_. Plasmids were transfected using the PEI MAX transfection reagent (Polysciences, Inc., Warrington, PA, USA), and protein expression was analyzed 48–72 hours post-transfection.

### 4.11. Plasmid construction

Full-length mouse Twf1 (mTwf1) was amplified using RT-PCR from mouse neurons. Total RNA was extracted using the RNeasy Mini Kit (QIAGEN, Hilden, Germany) according to the manufacturer’s instructions. PCR products were cloned into the pCRII-Blunt TOPO vector (Invitrogen, Eugene, OR, USA). The cloning primers used were as follows: mTWF1 1F, 5′-ATGTCCCACCAGACAGGCATCC-3′; mTWF1 1053R(+), 5′-CTAGTCAGTAGTGGCTTCTGCTTC-3′. After sequence confirmation, cDNA was subcloned into pCAGGS-Myc and pRSET-C1 vectors at the EcoRI site. Plasmids containing Src-GFP and Fyn-GFP were previously described [88]. To generate mutant Twf1, PCR was performed using a site-directed mutagenesis kit (Takara, Otsu, Japan). The primers used are listed in Table S3. Single-stranded oligonucleotides containing the target sequence for Twf1 were annealed to the pLLX vector (a kind gift from Dr. Michael Greenberg).

We used a modified pIRESpuro3 vector (Takara Bio, Mountain View, CA, USA) encoding a Flag sequence as a backbone plasmid. The plasmid was PCR-amplified using the 55/56 primer set to produce a linear DNA fragment. The hTWF1 cDNA, which was PCR-amplified from total RNA extracted from HEK293T cells using the 80/81 primer set, was inserted into a linearized plasmid using the InFusion HD cloning kit (Takara Bio, Tokyo, Japan), yielding hTWsF1 WT-Flag. For plasmids co-expressing hTWF1-Flag and Myc-Src, the Myc sequence was inserted into hTWF1 WT-Flag using the 96/105 primer set. The plasmid was linearized using the 63/106 primer set, and human Src, which was PCR-amplified using the 97/98 primer set, was inserted using the same approach, resulting in hTWF1 WT-Flag+Myc-Src WT.

Twf1 Y309F was generated via site-directed mutagenesis using the 88/89 primer set. Similarly, the K298M (kinase-dead) and Y530F (constitutively active) mutations of Src were generated using the 113/114 and 111/112 primer sets, respectively. The primers used are listed in Table S3.

### 4.12. Recombinant reelin production

Recombinant reelin was produced as previously described [35]. Expression constructs for WT reelin with an N-terminal PA tag and the K2360/2467A (K2A) reelin mutant (a mutant unable to bind to the reelin receptor) were a kind gift from Dr. J. Takagi (Osaka University). To generate the K2A reelin construct, K2A reelin was subcloned into the SalI sites of phCMV3-PA-WT reelin, yielding phCMV3-PA-K2A reelin. Expi293F cells (Thermo Fisher Scientific) were transfected with either the WT reelin construct or the K2A reelin construct, using the Expi293F Expression System (Thermo Fisher Scientific). Four days post-transfection, the culture medium was collected and centrifuged at 3,000 × *g* for 5 minutes to remove cells and debris. Recombinant reelin was purified by incubating the supernatant with anti-PA tag antibody-coated beads (Fujifilm, Osaka, Japan) and eluted using a PA tag peptide. The eluted protein was concentrated approximately 10-fold using an Amicon Ultra Concentrator (Merck). The concentration of purified recombinant reelin was determined using SDS-PAGE and Coomassie Brilliant Blue staining, with bovine serum albumin as a standard.

### 4.13. Primary neuronal culture, recombinant reelin stimulation, and PP2 treatment

Primary cortical neurons were cultured as previously described [32,35,90], with minor modifications. Briefly, cortical tissues were extracted from pregnant mice at embryonic day (E) 15–16 and digested in Hank’s balanced salt solution (Gibco) containing 0.25% trypsin (Thermo Fisher Scientific) and 0.01% DNase I (Roche Diagnostics GmbH, Mannheim, Germany) at 37 °C for 15 minutes. The tissues were then mechanically dissociated by gentle pipetting, and the resulting cell suspension was filtered through a 70-μm cell strainer. Neurons were seeded at a density of 1 × 10^6^ cells/well on poly-D-lysine-coated 6-well plates (IWAKI, Tokyo, Japan) at DIV0. Cultured neurons were maintained in Neurobasal medium (Gibco) supplemented with B27 (Thermo Fisher Scientific) and 0.5 mM GlutaMAX (Thermo Fisher Scientific). Half of the conditioned medium was replaced with fresh medium every 3–4 days. For reelin stimulation, neurons were treated with 30 nM reelin-conditioned medium or control supernatant for 15 minutes. For SFK inhibition, PP2 (MedChemExpress, Monmouth Junction, NJ, USA; HY-13805) was dissolved in dimethyl sulfoxide (DMSO) to prepare a 200 mM stock solution, which was stored at −20 °C. The stock solution was diluted in culture medium to a final concentration of 10 μM, ensuring that the DMSO concentration remained below 0.1%. Cultured neurons were pre-treated with PP2 for 1 hour before reelin stimulation.

### 4.14. Lentivirus production, quantification, and microinjection

Lentivirus production and quantification were performed as previously described [76], Briefly, HEK293T cells were transfected with a transfer plasmid, envelope plasmid, and packaging plasmid in Opti-MEM (Gibco) using PEI MAX transfection reagent (Polysciences, Inc.). After overnight incubation at 37 °C, the medium was replaced with fresh culture medium. Virus-containing supernatants were collected 24 hours after medium replacement, centrifuged at 3,000 × *g* for 10 minutes, and further concentrated by centrifugation at 6,000 × *g* for 16–18 hours. The lentivirus-containing pellet was resuspended in 200 μL of Dulbecco’s phosphate-buffered saline (Gibco). For lentivirus copy number quantification, total RNA was extracted using the QIAamp Viral RNA Mini Kit (QIAGEN), and RNA concentrations were measured using a NanoDrop 2000c spectrophotometer (Thermo Fisher Scientific). cDNA was synthesized from purified RNA using the Superscript III First-Strand Synthesis SuperMix for qRT-PCR (Invitrogen). qRT-PCR was conducted as described above. Data were analyzed using a standard curve generated from a pLLX plasmid containing EGFP, and relative quantification was performed using the 2^−ΔΔCT^ method. The lentivirus was diluted to a final concentration of 10^8^ transduction units/mL and stored at −80 °C in 10-μL aliquots.

Microinjection was performed as previously described [35], with minor modifications. Mice were first anesthetized with tribromoethanol (Avertin; 250 mg/kg, i.p.) and positioned in a stereotaxic frame (David Kopf Instruments, Tujunga, CA, USA). A small hole was drilled through the skull to allow direct needle passage into the bilateral mPFC. A total of 0.5 μL of lentivirus was injected into the bilateral mPFC at the following stereotaxic coordinates: +1.5 mm anteroposterior, ±0.5 mm mediolateral, and −2.5 mm dorsoventral from the bregma. Mice were left for three weeks post-injection to allow for sufficient infection and transgene expression.

### 4.15. Immunohistochemistry

Immunohistochemistry was performed as previously described [91], with minor modifications. Mice were anesthetized with tribromoethanol (250 mg/kg, i.p.) and transcardially perfused with ice-cold 0.1 M phosphate buffer (PB) followed by 4% paraformaldehyde (PFA) in PB. The brains were extracted and post-fixed in 4% PFA, then cryoprotected by sequential incubation in 20% and 30% sucrose in PB. Frozen brain sections (30 μm thick) were cut coronally using a cryostat (Leica). Cryosections were fixed with 4% PFA in 0.1 M PB for 5 minutes and then permeabilized with 0.3% Triton X-100 in PBS for 10 minutes. After blocking with 5% goat serum (Vector Laboratories Inc.) in PBS containing 0.3% Triton X-100 for 1 hour at room temperature, sections were incubated with primary antibodies overnight at 4 °C. The next day, sections were washed three times with PBS and incubated with secondary antibodies at room temperature for 1 hour. Following PBS washes, sections were mounted on adhesive silane (MAS)–coated glass slides (Matsunami Glass Ind. Ltd., Osaka, Japan) using Fluorescent Mounting Medium (Dako, Santa Clara, CA, USA) and a coverslip. Images of dendrites were acquired using a TiR-A1R confocal laser microscope (Nikon) with a 100× objective lens under oil immersion. All imaging data were analyzed by an investigator blinded to the genotypes of the mouse brains. Antibodies used in immunohistochemistry are listed in Table S2.

### 4.16. Spine analysis

Dendrites and spines located 50–200 µm from the soma in the secondary or tertiary branches of pyramidal neurons were analyzed, with 6–7 dendrites per mouse examined. Z-stacks with 0.2 μm step intervals were acquired to image spines, and 3D reconstructions were generated from fluorescence images using Imaris software version 9.5 (Oxford Instruments). Spine density was determined by counting the number of spines per 10 µm of dendrite length. According to the morphological classification rules provided by Oxford Instruments, protrusions were categorized into four types: (1) mushroom spines with a large head (maximum head width□> 2 × mean neck width, spine length ≥ 1 μm); (2) stubby spines with a head but no neck (spine length < 1 μm); (3) thin spines with a long neck and a small head (mean head width ≥ mean neck width, maximum head width ≤ 2 × mean neck width, spine length ≥ 1 μm); and (4) filopodia spines with a long neck but no head (mean neck width > mean□head width, spine length ≥ 1 μm).

### 4.17. Twf1 subcellular localization

Cortical neurons were seeded at DIV0 at a density of 5□×□10^5^ cells/well on poly-D-lysine-coated 27-mm glass-based dishes (IWAKI). At DIV15, Myc-tagged Twf1 WT, Twf1 Y309E, or Twf1 Y309F constructs were co-transfected with Syn-mCherry into neurons using Lipofectamine 2000 (Invitrogen). Three days after transfection, cortical neurons were fixed with freshly prepared 4% PFA in 0.1 M PB for 15 minutes, followed by permeabilization with 0.1% Triton X-100 for 15 minutes. After blocking in 5% goat serum in PBS for 1 hour at room temperature, neurons were incubated with primary antibodies diluted in blocking buffer at 4 °C overnight. The next day, neurons were washed three times with PBS and incubated with secondary antibodies at room temperature for 1 hour. Confocal images were captured using a TiE-A1R confocal laser scanning microscope (Nikon). Spines were imaged by acquiring Z-stacks with 0.2-μm step intervals, and 2D images were created using maximum intensity projections. Fluorescence intensity was quantified using Fiji software, with all samples blinded before image analysis. The spine-to-shaft ratio [43] was determined by measuring the average fluorescence intensity of Myc in the spine head and comparing it to that of an equal-sized area in the adjacent dendrite. Measurements were normalized to the intensity distribution of mCherry within the corresponding regions.

### 4.18. Layer thickness measurement

Coronal brain sections from 8-week-old WT, heterozygous *Twf1^+/Y309F^*, and homozygous *Twf1^Y309F/Y309F^* mice were subjected to immunohistochemistry as described in the main text. Anti-Cux1 and anti-Ctip2 antibodies were used as markers for specific cortical layers [48,49]. Fluorescent images of cortical layers were captured using a BZ-X800 microscope (Keyence) equipped with a 10× objective lens. In the adult mouse brain, Cux1-positive cells are typically located in layers II–III, while Ctip2-positive cells are found in layers V–VI. The thickness of Cux1-positive and Ctip2-positive cell layers was measured, along with the distance between these layers. Measurements were obtained from multiple coronal brain sections at three defined points in each cortical hemisphere. Cortical thickness was quantified using the BZ-X800 Analyzer (Keyence).

### 4.19. F/G-actin assay

The levels of polymerized F-actin and monomeric G-actin in HEK293T cells and cultured neurons were quantified using the G-actin/F-actin In Vivo Assay Kit (Cytoskeleton, Inc., Denver, CO, USA; BK103), following the manufacturer’s instructions. Briefly, cells were lysed in pre-warmed lysis and F-actin stabilizing buffer, supplemented with protease inhibitor and 1 mM ATP. The cell lysates were incubated at 37 °C for 10 minutes and then centrifuged for 5 minutes at 350 × *g* to remove cell debris. The lysates were ultracentrifuged at 100,000 × *g* for 1 hour at 37 °C to pellet F-actin, leaving G-actin in the supernatant. The supernatants were transferred to new tubes and designated as G-actin samples. F-actin pellets were resuspended in F-actin depolymerization buffer and incubated on ice for 1 hour with periodic pipetting to ensure complete depolymerization. Equal volumes of G-actin and F-actin fractions were then mixed with 5x SDS sample buffer and analyzed via western blot using the anti-actin antibody provided in the kit.

### 4.20. Behavioral tests

All mice were maintained in a specific pathogen-free environment. Mice were housed in groups of 4–6 under standard conditions (23 ± 1 °C, 50 ± 5% humidity) with a 12-hour light/dark cycle (09:00–21:00 light phase). All experiments were performed during the light phase. Food and water were provided *ad libitum* throughout the experiments, except during the touchscreen-based VD and RL tasks. Male and female mice aged 8–20 weeks were used in the experiments. Behavioral experiments were conducted in a sound-attenuated, air-regulated room where mice were habituated for at least 1 hour before testing. Behavioral tests were conducted in the following order: locomotor activity, open field, Y-maze, social interaction, and touchscreen-based VD and RL tasks. The protocols used were described previously [50].

#### 4.20.1. Locomotor activity test

Locomotor activity was assessed by placing each mouse individually in a transparent rectangular cage (25 × 25 × 20 cm) with a black frosted Plexiglas floor under moderate illumination (15 lx). Locomotor activity was recorded every 5 minutes for 120 minutes using infrared sensor-equipped digital counters (BrainScience Idea, Osaka, Japan).

#### 4.20.2 Open field test

Mice were placed in the center of a circular open field (diameter: 60 cm, height: 35 cm) under moderate light conditions (85 lx) and allowed to explore freely for 5 minutes. Movements of mice were recorded using the EthoVision automated tracking program (BrainScience Idea Co., Ltd., Osaka, Japan) with an overhead camera. The open field was divided into an inner circle (diameter: 40 cm) and an outer area. Measured parameters included time spent in each zone and total distance traveled. Additionally, rearing, jumping, grooming, defecation, and urination events were manually counted.

#### 4.20.3. Y-maze test

The Y-maze test was performed as previously described [90,92]. Mice were placed at the center of a Y-maze (arm dimensions: 40 cm long, 12 cm high, 3 cm wide at the bottom, and 10 cm wide at the top; arms set 120° apart) and allowed to move freely through the maze for 8 minutes under 35 lx illumination. Arm entries were recorded visually. Spontaneous alternation, defined as consecutive entries into all three arms in overlapping triplet sequences (e.g., ABC, CAB, BCA, but not BBC), was used to assess short-term memory capacity. Percent alternation was calculated as the ratio of actual alternations to the total possible alternations, where the latter can be determined by subtracting two from the total number of arm entries. This was expressed using the following equation: % alternation = (number of alternations)/(total arm entries – 2) × 100.

#### 4.20.4. Social interaction test

The social interaction test was conducted as described previously [93] with minor modifications. The apparatus consisted of a black rectangular open arena (30 × 30 × 35 cm) without a top cover. All sessions were conducted under low illumination (15 lx). Mice were habituated to the test box for 10 minutes per day over two consecutive days prior to testing. On the test day, each test mouse was randomly assigned to an unfamiliar sex-, age-, and strain-matched stimulus mouse. Both the test mouse and the stimulus mouse were placed in the test box for 10 minutes while their social behaviors were observed. The total duration of social interactions, including sniffing, grooming, following, mounting, and crawling (excluding aggressive behaviors), was recorded.

### 4.21. Touchscreen-based VD and RL tasks

The protocol used in these tasks was previously described [45,94]. Briefly, 8-week-old mice were restricted to food and water access to a 2-hour window (17:00–19:00) each day for 1 week before pretraining to ensure sufficient motivation to perform the tasks. This restriction was maintained throughout the task, ensuring that the mice’s body weight remained at 85–90% of that of unrestricted animals. Next, mice underwent pretraining, which consisted of five stages—habituation, initial touch, must touch, must initiate, and punish incorrect—to shape screen-touching behavior. After mice learned to operate the touchscreen (≥75% correct responses for two consecutive days), they proceeded to the VD and RL tasks.

#### 4.21.1. VD task

Each trial was initiated when mice touched the nozzle, after which two stimuli (fan and marble) were randomly presented in two response windows simultaneously. Touching the correct window resulted in a reward (20 μL of milk), whereas touching the incorrect window triggered an immediate stimulus offset and a 5-second time-out punishment period. After a 20-second inter-trial interval, mice received a correction trial instead of a new trial. During the correction trial, the same stimuli were repeatedly presented in the same response window until mice made a correct response. The stimulus contingencies were counterbalanced. Each session ended after 30 trials. Mice were considered to have completed the VD task once they achieved a correct response rate of ≥80% for two consecutive days.

#### 4.21.2. RL task

Once mice met the VD task criterion, they proceeded to the RL task. The RL task closely resembled the initial acquisition phase of the VD task, with the key difference being the reversal of stimulus contingencies. The stimulus that was previously rewarded was now designated as incorrect, whereas the previously non-rewarded stimulus became the correct choice.

### 4.22. Neuronal induction

The iPSCs derived from the peripheral blood of three individuals (two healthy donors and one patient with schizophrenia (harboring a *RELN*-del) were differentiated into neurospheres, following a previously reported protocol [53,82,95], with minor modifications. Briefly, after passaging onto feeder cells, the iPSCs were treated with SB431542, CHIR99021, and dorsomorphin. After 7 days, they were dissociated into single cells and cultured in ultra-low attachment 6-well plates for 2 weeks. Neurospheres were passaged by dissociating them into single cells.

### 4.23. Filopodia analysis

Neurospheres were dissociated into single cells using Accutase and plated onto poly-L-ornithine-coated 8-well chambers. After 24 hours, cells were fixed in 4% PFA for 15 minutes. F-actin was stained using Alexa Fluor 488-Phalloidin (Cytoskeleton), followed by DAPI staining, as previously described [96]. Filopodia morphology was analyzed using the open-source software FiloQuant [97].

### 4.24. Statistical analysis

All data are presented as mean ± standard error of the mean. Statistical analyses were conducted using GraphPad Prism version 9 (GraphPad Inc., San Diego, CA, USA). Statistical significance (P < 0.05) was determined using one-way analysis of variance, mixed-effects analysis, unpaired Student’s *t*-test, log-rank test, or the Kolmogorov–Smirnov test for multigroup comparisons. Tukey’s, Dunnett’s, or Šídák’s multiple comparisons tests were applied for post-hoc analysis. Detailed information on statistical analyses is provided in Table S4.

## Abbreviations

Twf1: Twinfilin-1
ADF-H: actin-depolymerizing factor homology
RELN: reelin
ApoER2: apolipoprotein E receptor 2
VLDLR: very low-density lipoprotein receptor
CP: capping protein
Dab1: Disabled homolog 1
DIV: days in vitro
SFKs: Src family kinases
Src: proto-oncogene tyrosine-protein kinase Src
Fyn: proto-oncogene tyrosine-protein kinase Fyn
HEK293T: human embryonic kidney 293T cells
hTWF1: human Twinfilin-1
iPSC: induced pluripotent stem cell
CA: constitutively active
KD: kinase-dead
LC-MS/MS: liquid chromatography–tandem mass spectrometry
mPFC: medial prefrontal cortex
pTwf1: phosphorylated Twf1
pY: phosphotyrosine
PFA: paraformaldehyde
VD: visual discrimination
RL: reversal learning
WT: wild-type
Y309: tyrosine 309.

## CRediT authorship contribution statement

**Geyao Dong:** Conceptualization, Methodology, Investigation, Writing – original draft, Writing – review & editing. **Daisuke Mori:** Methodology, Investigation, Writing – review & editing. **Tetsuo Matsuzaki:** Methodology, Investigation, Writing – review & editing. **Rinako Tanaka:** Methodology, Investigation, Writing – review & editing. **Norimichi Itoh:** Methodology, Investigation, Writing – review & editing. **Takaaki Matsui:** Methodology, Investigation, Writing – review & editing. **Ayato Sato:** Methodology, Investigation, Writing – review & editing. **Yuko Arioka:** Methodology, Investigation, Writing – review & editing. **Hiroki Okumura:** Methodology, Investigation, Writing – review & editing. **Ryota Fukaya:** Methodology, Investigation, Writing – review & editing. **Hiroshi Kuba:** Writing – review & editing, Supervision. **Taku Nagai:** Writing – review & editing, Supervision. **Toshitaka Nabeshima:** Writing – review & editing, Supervision. **Hiroaki Ikesue:** Writing – review & editing, Supervision. **Takao Kohno:** Methodology, Investigation, Writing – review & editing. **Mitsuharu Hattori:** Writing – review & editing, Supervision. **Kozo Kaibuchi:** Writing – review & editing, Supervision. **Norio Ozaki:** Writing – review & editing, Supervision. **Hiroyuki Mizoguchi:** Conceptualization, Methodology, Writing – review & editing, Supervision, Funding acquisition. **Kiyofumi Yamada:** Conceptualization, Methodology, Writing – review & editing, Supervision, Funding acquisition.

## Ethical approval and consent

Generation and application of human iPSCs were approved by the Nagoya University Ethics Committee (approval number: 2012–0184). The procurement of human iPSCs was conducted in accordance with the WHO *Guiding Principles on Human Cell, Tissue, and Organ Transplantation*. The study conforms to the principles outlined in the Declaration of Helsinki. All subjects provided written informed consent.

All procedures involving animals adhered to the guidelines established by the Institutional Animal Care and Use Committee of Nagoya University, the *Guiding Principles for the Care and Use of Laboratory Animals* set forth by the Japanese Pharmacological Society, and the National Institutes of Health *Guide for the Care and Use of Laboratory Animals*.

## Acknowledgements

We thank Dr. J. Takagi (Osaka University) for providing the phCMV3-K2A reelin plasmid. We are also grateful to the staff (Eri Yorifugi, Mayumi Furukawa, Yasutomo Ito, Moeko Marui, and Kentaro Taki) of the Division for Medical Research Engineering at the Nagoya University Graduate School of Medicine for their assistance with the BZ-X800 microscope and analyzer (Keyence), TiR-A1R confocal microscope (Nikon), NIS-Elements analysis software (Nikon), Imaris software (Oxford Instruments), and the Optima MAX-XP ultracentrifuge (Beckman Coulter). We appreciate their technical support and assistance with data acquisition. G.D. acknowledges the support of the Kobayashi Scholarship Foundation.

## Appendix A. Supporting information

Supplementary data associated with this article can be found in the online version at [link].

## Data Availability

Data will be made available on request.

## Supplementary material

**Fig. S1.**
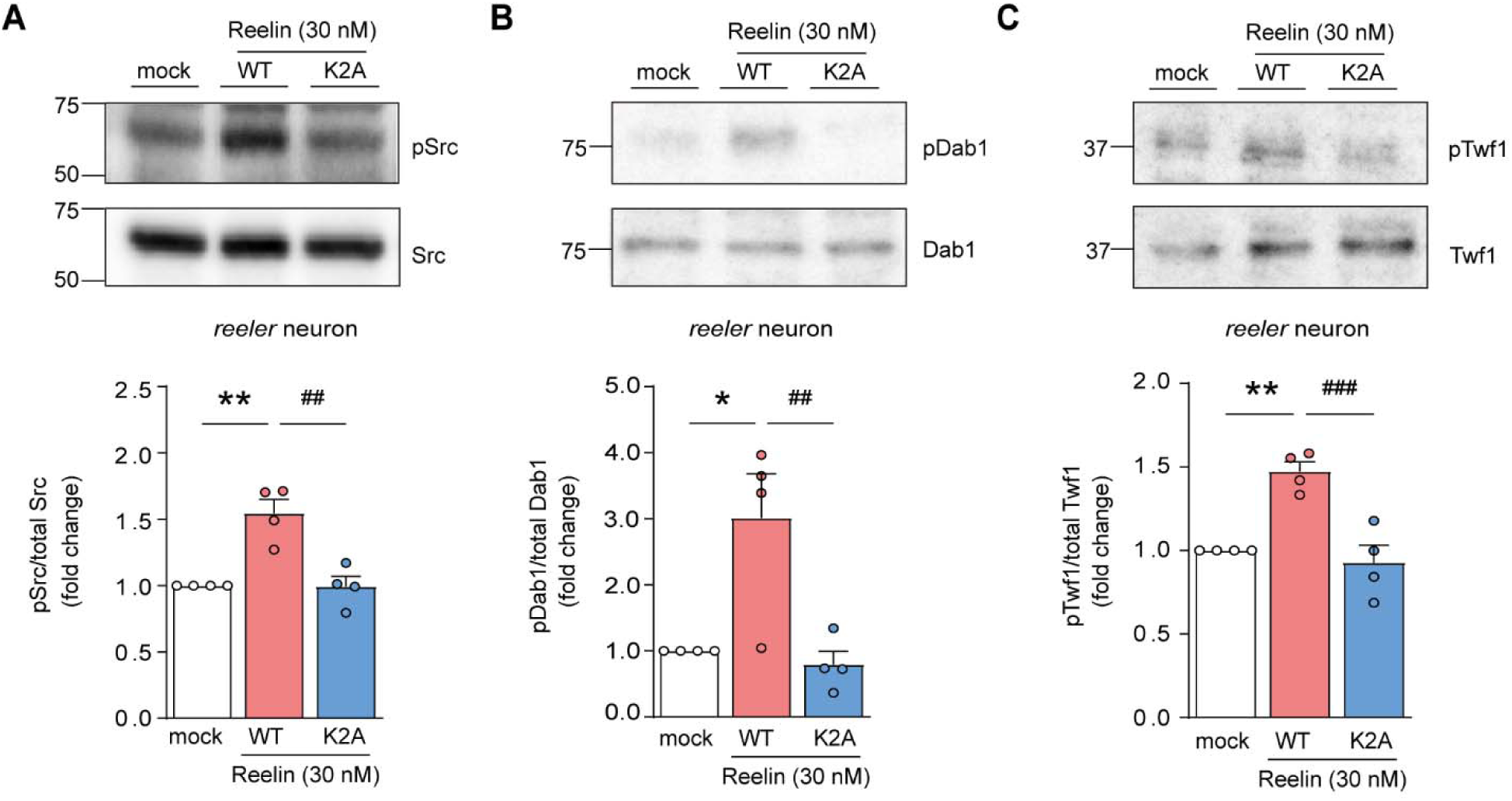
Recombinant reelin induces Twf1 phosphorylation in primary cortical neurons from reeler mice. (A–C) Effects of WT and K2A reelin on the phosphorylation levels of (A) Src, (B) Dab1, and (C) Twf1 in primary cultured cortical neurons from *reeler* mice. Cultured neurons were treated with WT or K2A reelin at 30 nM on DIV5 for 15 min. ** P < 0.01, * P < 0.05 vs. mock group; ^###^ P < 0.001, ^##^ P < 0.01 vs. WT reelin group, Tukey’s multiple comparisons test (n = 3 independent experiments). All data represent the mean ± SEM.

**Fig. S2.**
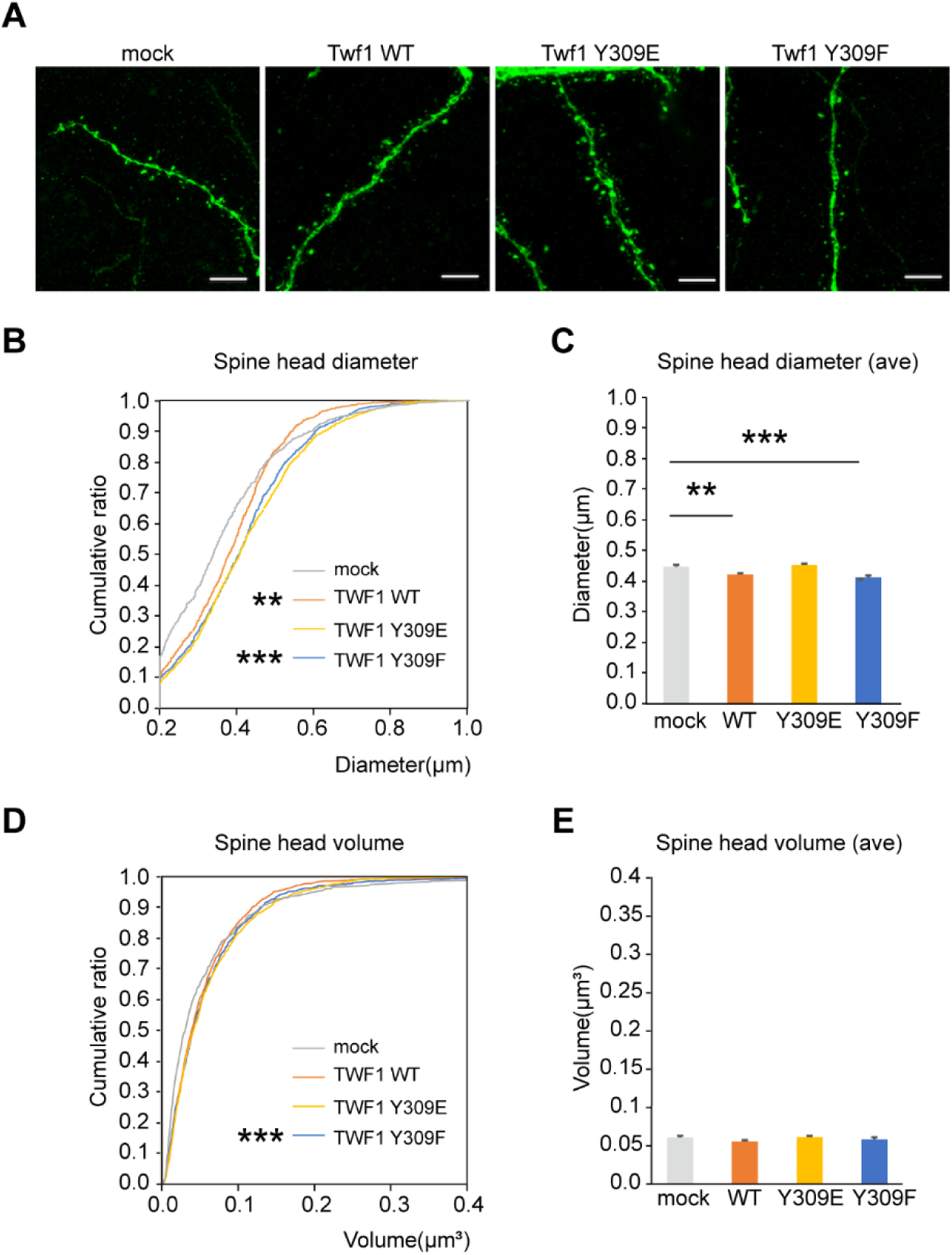
Expression of phospho-resistant Twf1 Y309F in the mPFC of mice reduces dendritic spine head size. (A) Representative images of dendritic spines in the mPFC of WT mice injected with lentivirus encoding GFP, WT Twf1-GFP, phospho-mimic Twf1 Y309E-GFP, or phospho-resistant Twf1 Y309F-GFP. Scale bar: 10 µm. (B and C) Effects of Twf1 phosphorylation on spine head diameter. (B) Cumulative ratio, *** P < 0.001, ** P < 0.01 vs. mock group, Kolmogorov–Smirnov test. c, Average diameter, *** P < 0.001, ** P < 0.01 vs. mock group, Tukey’s multiple comparisons test. (D and E) Effect of Twf1 phosphorylation on spine head volume. (D) Cumulative ratio, *** P < 0.001 vs. mock group, Kolmogorov–Smirnov test. (E) Average volume. n = 778–1405 spines from 4–6 mice. All data represent the mean ± SEM.

**Fig. S3.**
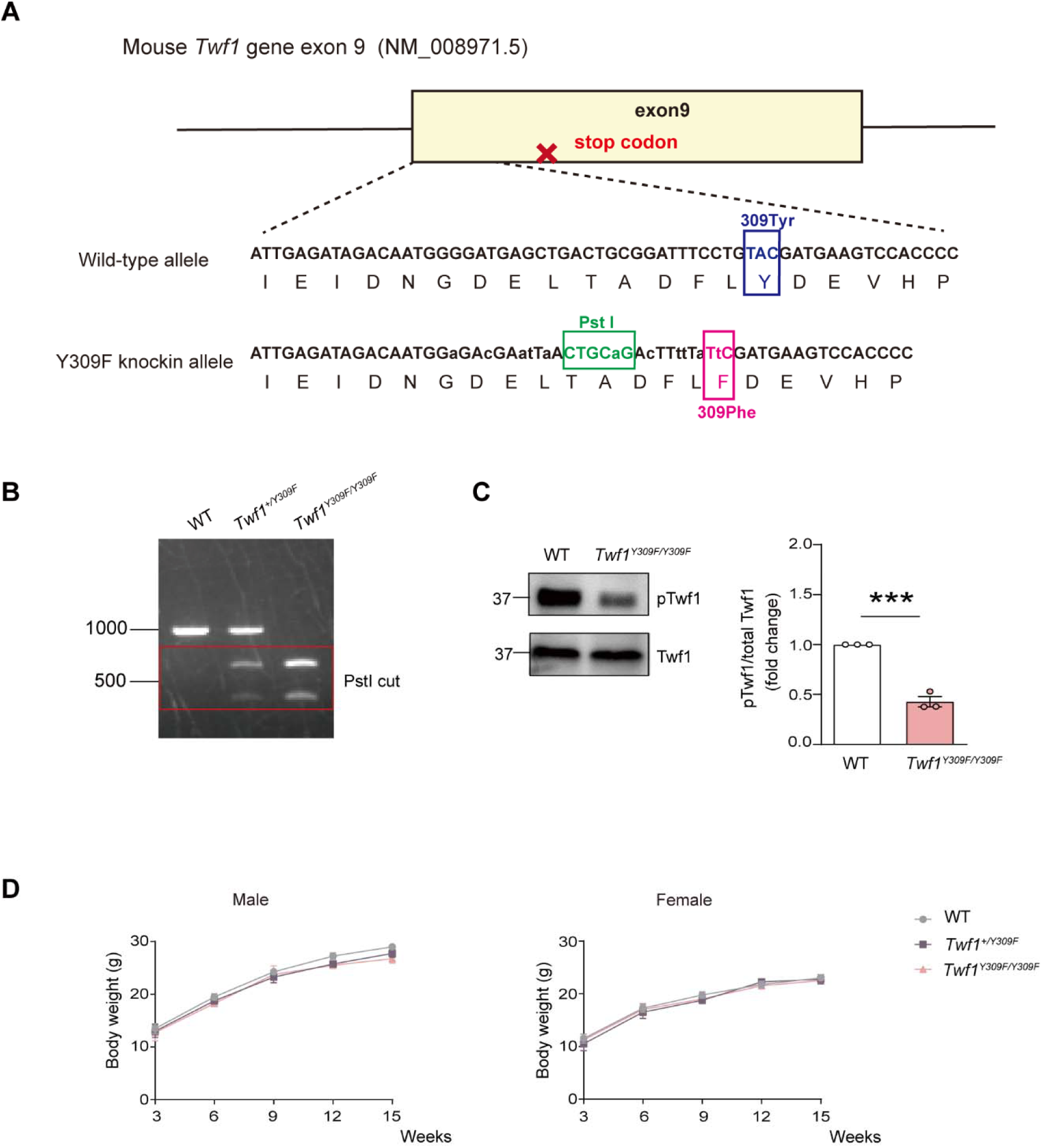
Generation and characterization of Twf1^Y309F^ mice. (A) Schematic representation of *Twf1^Y309F^*mouse generation using CRISPR/Cas9 gene editing. (B) PCR genotyping of WT, heterozygous *Twf1^+/Y309F^*, and homozygous *Twf1^Y309F/Y309F^*mice. Mutant alleles exhibit dual bands cleaved by PstI. (C) Twf1 phosphorylation levels in the brains of WT and homozygous *Twf1^Y309F/Y309F^*mice. *** P < 0.001 vs. WT, unpaired Student’s *t*-test (n = 3 mice/group). (D) Body weight measurements of WT, heterozygous *Twf1^+/Y309F^*, and homozygous *Twf1^Y309F/Y309F^*mice from 3–15 weeks. Mixed-effects analysis, Šídák’s multiple comparisons test (n = 4 mice/group). All data represent the mean ± SEM.

**Fig. S4.**
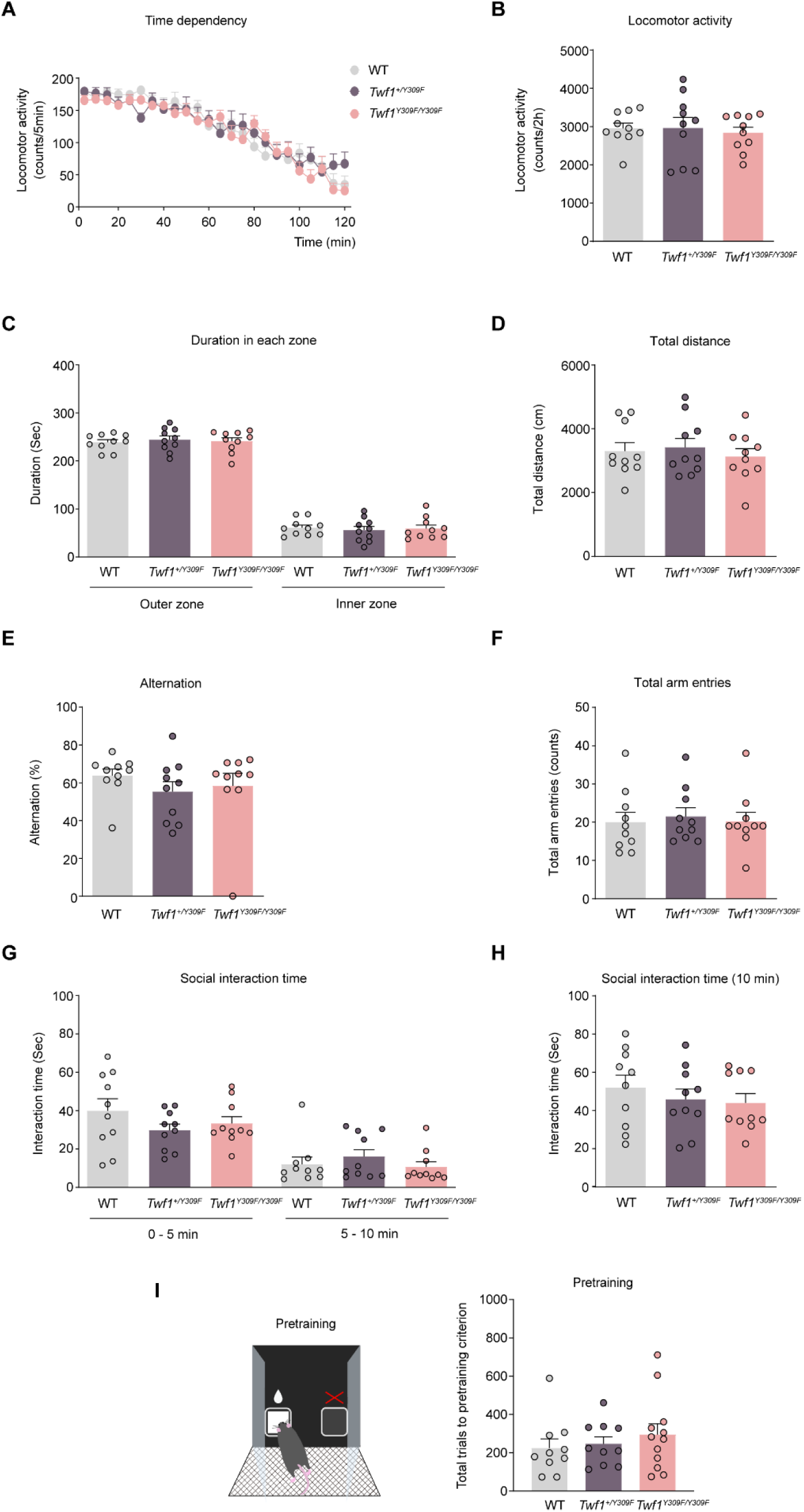
Behavioral analyses in Twf1^Y309F^ mice. Performance of WT, heterozygous *Twf1^+/Y309F^*, and homozygous *Twf1^Y309F/Y309F^* mice in the locomotor activity test (A and B), open field test (C and D), Y-maze test (E and F), social interaction test (G and H), and pre-training for touchscreen-based behavioral tasks (I). WT mice (n = 10, male = 5, female = 5), *Twf1^+/Y309F^* mice (n = 10, male = 4, female = 6), and *Twf1^Y309F/Y309F^* mice (n = 10, male = 5, female = 5) for (A–H). WT mice (n = 10, male = 5, female = 5), *Twf1^+/Y309F^*mice (n = 10, male = 5, female = 5), and *Twf1^Y309F/Y309F^* mice (n = 12, male = 6, female = 6) for (I). Two-way repeated measures analysis of variance (ANOVA) followed by Dunnett’s multiple comparisons test was used for A. Two-way ANOVA followed by Dunnett’s multiple comparisons test (C and G). One-way ANOVA followed by Dunnett’s multiple comparisons test (B, D, E, F, H, and I). All data represent the mean ± SEM.

**Fig. S5.**
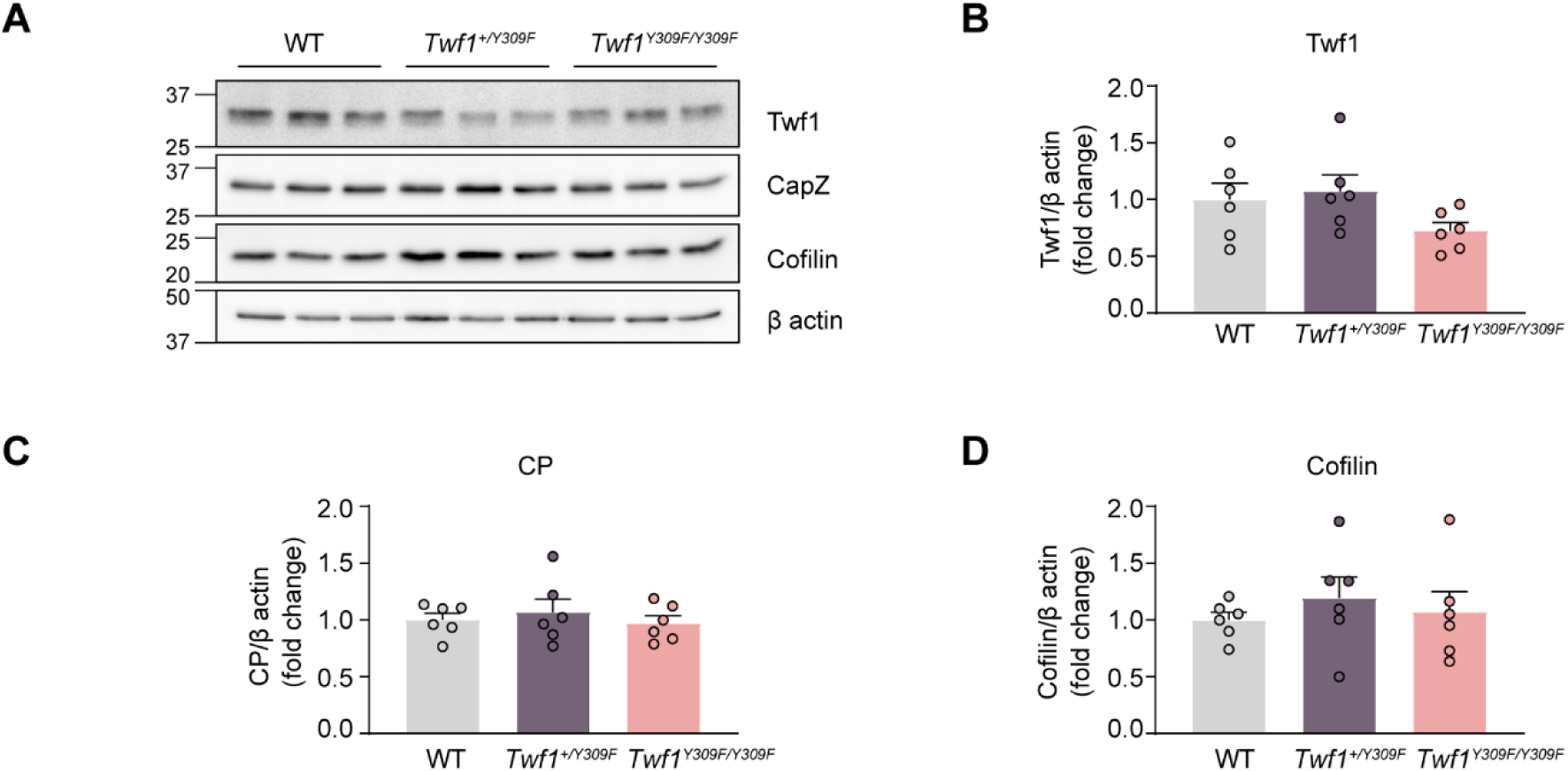
Twf1^Y309F^ mice exhibit no changes in levels of Twf1-associated proteins. (A) Representative western blots showing Twf1, CP, and cofilin levels in the mPFC of WT, heterozygous *Twf1^+/Y309F^*, and homozygous *Twf1^Y309F/Y309F^*mice. (B–D) Quantification of Twf1 (B), CP (C), and cofilin (D) levels. n = 6 mice/group (male = 3, female = 3). One-way ANOVA followed by Tukey’s multiple comparisons test. All data represent the mean ± SEM.

**Fig. S6.**
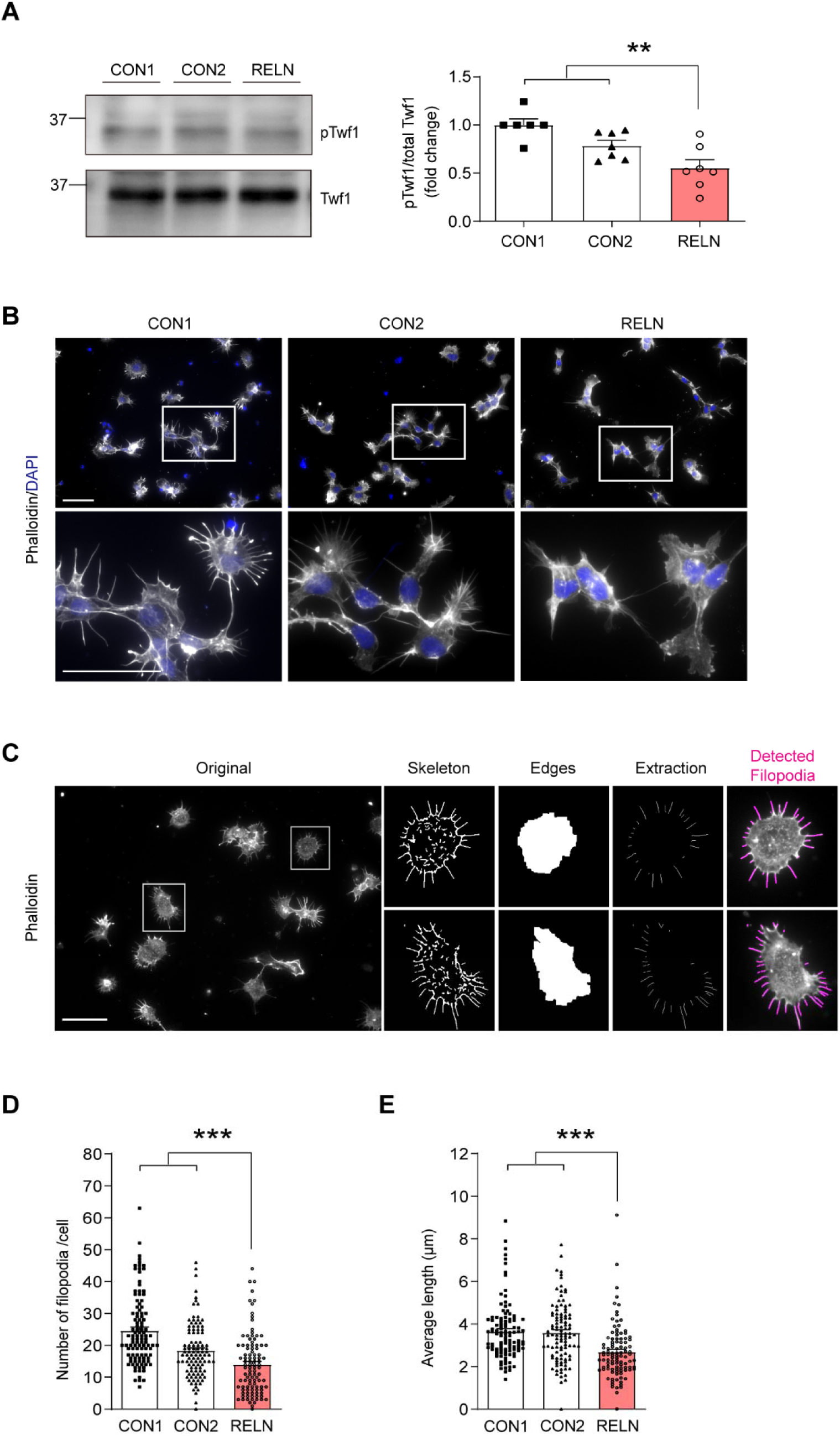
RELN-del iPSCs from a patient with schizophrenia exhibit reduced Twf1 phosphorylation and disrupted F-actin dynamics. (A) Twf1 phosphorylation levels in *RELN*-del iPSCs were analyzed by immunoprecipitation using an anti-pY antibody, followed by western blotting with an anti-Twf1 antibody. ** P < 0.01 vs. CON group, unpaired Student’s *t*-test. CON group: n = 13 (CON1: n = 6; CON2: n = 7); RELN group: n = 7. (B) Representative images of CON1, CON2, and *RELN*-del iPSC-derived neurospheres stained with phalloidin and DAPI. Scale bar: 50 µm. c, Representative FiloQuant analysis results, with detected filopodia highlighted in magenta in the far-right images. Scale bar: 50 µm. (D and E) Analysis of F-actin dynamics in iPSC-derived neurospheres, showing the number of filopodia per cell (D) and the average length of filopodia (E). *** P < 0.001 vs. CON group, unpaired Student’s *t*-test. CON group: n = 212 (CON1: n = 108; CON2: n = 104); RELN group: n = 103. All data represent the mean ± SEM.

## References

[1] G. D’Arcangelo, G.G. Miao, S.C. Chen, H.D. Soares, J.I. Morgan, & T. Curran. A protein related to extracellular matrix proteins deleted in the mouse mutant reeler. Nature. 374 (1995) 719–723. 10.1038/374719a0.

[2] F. Tissir & A.M. Goffinet. Reelin and brain development. Nat. Rev. Neurosci. 4 (2003) 496–505. 10.1038/nrn1113.

[3] C. Pesold, F. Impagnatiello, M.G. Pisu, D.P. Uzunov, E. Costa, A. Guidotti, & H.J. Caruncho. Reelin is preferentially expressed in neurons synthesizing γ-aminobutyric acid in cortex and hippocampus of adult rats. Proc. Natl. Acad. Sci. U. S. A. 95 (1998) 3221–3226. 10.1073/pnas.95.6.3221.

[4] Y. Jossin. Reelin Functions, Mechanisms of Action and Signaling Pathways During Brain Development and Maturation. Biomolecules. 10 (2020) 964. 10.3390/biom10060964.

[5] S. Niu, O. Yabut, & G. D’Arcangelo. The Reelin signaling pathway promotes dendritic spine development in hippocampal neurons. J. Neurosci. 28 (2008) 10339–10348. 10.1523/jneurosci.1917-08.2008.

[6] Z. Zhou, Z. Hu, L. Zhang, Z. Hu, H. Liu, Z. Liu, J. Du, J. Zhao, L. Zhou, K. Xia, B. Tang, & L. Shen. Identification of RELN variation p.Thr3192Ser in a Chinese family with schizophrenia. Sci. Rep. 6 (2016) 24327. 10.1038/srep24327.

[7] G. Costain, A.C. Lionel, D. Merico, P. Forsythe, K. Russell, C. Lowther, T. Yuen, J. Husted, D.J. Stavropoulos, M. Speevak, E.W. Chow, C.R. Marshall, S.W. Scherer, & A.S. Bassett. Pathogenic rare copy number variants in community-based schizophrenia suggest a potential role for clinical microarrays. Hum. Mol. Genet. 22 (2013) 4485–4501. 10.1093/hmg/ddt297.

[8] A. Sobue, I. Kushima, T. Nagai, W. Shan, T. Kohno, B. Aleksic, Y. Aoyama, D. Mori, Y. Arioka, N. Kawano, M. Yamamoto, M. Hattori, T. Nabeshima, K. Yamada, & N. Ozaki. Genetic and animal model analyses reveal the pathogenic role of a novel deletion of RELN in schizophrenia. Sci. Rep. 8 (2018) 13046. 10.1038/s41598-018-31390-w.

[9] I. Kushima, B. Aleksic, M. Nakatochi, T. Shimamura, T. Shiino, A. Yoshimi, H. Kimura, Y. Takasaki, C. Wang, J. Xing, K. Ishizuka, T. Oya-Ito, Y. Nakamura, Y. Arioka, T. Maeda, M. Yamamoto, M. Yoshida, H. Noma, S. Hamada, M. Morikawa, Y. Uno, T. Okada, T. Iidaka, S. Iritani, T. Yamamoto, M. Miyashita, A. Kobori, M. Arai, M. Itokawa, M.C. Cheng, Y.A. Chuang, C.H. Chen, M. Suzuki, T. Takahashi, R. Hashimoto, H. Yamamori, Y. Yasuda, Y. Watanabe, A. Nunokawa, T. Someya, M. Ikeda, T. Toyota, T. Yoshikawa, S. Numata, T. Ohmori, S. Kunimoto, D. Mori, N. Iwata, & N. Ozaki. High-resolution copy number variation analysis of schizophrenia in Japan. Mol. Psychiatry. 22 (2017) 430–440. 10.1038/mp.2016.88.

[10] F. Impagnatiello, A.R. Guidotti, C. Pesold, Y. Dwivedi, H. Caruncho, M.G. Pisu, D.P. Uzunov, N.R. Smalheiser, J.M. Davis, G.N. Pandey, G.D. Pappas, P. Tueting, R.P. Sharma, & E. Costa. A decrease of reelin expression as a putative vulnerability factor in schizophrenia. Proc. Natl. Acad. Sci. U. S. A. 95 (1998) 15718–15723. 10.1073/pnas.95.26.15718.

[11] A. Guidotti, J. Auta, J.M. Davis, V. Di-Giorgi-Gerevini, Y. Dwivedi, D.R. Grayson, F. Impagnatiello, G. Pandey, C. Pesold, R. Sharma, D. Uzunov, & E. Costa. Decrease in reelin and glutamic acid decarboxylase67 (GAD67) expression in schizophrenia and bipolar disorder: a postmortem brain study. Arch. Gen. Psychiatry. 57 (2000) 1061–1069. 10.1001/archpsyc.57.11.1061.

[12] S.H. Fatemi, J.M. Stary, J.A. Earle, M. Araghi-Niknam, & E. Eagan. GABAergic dysfunction in schizophrenia and mood disorders as reflected by decreased levels of glutamic acid decarboxylase 65 and 67 kDa and Reelin proteins in cerebellum. Schizophr. Res. 72 (2005) 109–122. 10.1016/j.schres.2004.02.017.

[13] S.L. Eastwood & P.J. Harrison. Cellular basis of reduced cortical reelin expression in schizophrenia. Am. J. Psychiatry. 163 (2006) 540–542. 10.1176/appi.ajp.163.3.540.

[14] G.G. D’Arcangelo, R. Homayouni, L. Keshvara, D.S. Rice, M. Sheldon, & T. Curran. Reelin is a ligand for lipoprotein receptors. Neuron. 24 (1999) 471–479. 10.1016/s0896-6273(00)80860-0.

[15] T. Hiesberger, M. Trommsdorff, B.W. Howell, A. Goffinet, M.C. Mumby, J.A. Cooper, & J. Herz. Direct binding of Reelin to VLDL receptor and ApoE receptor 2 induces tyrosine phosphorylation of disabled-1 and modulates tau phosphorylation. Neuron. 24 (1999) 481–489. 10.1016/s0896-6273(00)80861-2.

[16] B.W. Howell, T.M. Herrick, & J.A. Cooper. Reelin-induced tyrosine [corrected] phosphorylation of disabled 1 during neuronal positioning. Genes Dev. 13 (1999) 643–648. 10.1101/gad.13.6.643.

[17] B.W. Howell, T.M. Herrick, J.D. Hildebrand, Y. Zhang, & J.A. Cooper. Dab1 tyrosine phosphorylation sites relay positional signals during mouse brain development. Curr. Biol. 10 (2000) 877–885. 10.1016/s0960-9822(00)00608-4.

[18] L. Arnaud, B.A. Ballif, E. Förster, & J.A. Cooper. Fyn tyrosine kinase is a critical regulator of disabled-1 during brain development. Curr. Biol. 13 (2003) 9–17. 10.1016/s0960-9822(02)01397-0.

[19] H.H. Bock & J. Herz. Reelin activates SRC family tyrosine kinases in neurons. Curr. Biol. 13 (2003) 18–26. 10.1016/s0960-9822(02)01403-3.

[20] Y. Jossin & J.A. Cooper. Reelin, Rap1 and N-cadherin orient the migration of multipolar neurons in the developing neocortex. Nat. Neurosci. 14 (2011) 697–703. 10.1038/nn.2816.

[21] Y. Jossin & A.M. Goffinet. Reelin signals through phosphatidylinositol 3-kinase and Akt to control cortical development and through mTor to regulate dendritic growth. Mol. Cell. Biol. 27 (2007) 7113–7124. 10.1128/mcb.00928-07.

[22] X. Chai, E. Förster, S. Zhao, H.H. Bock, & M. Frotscher. Reelin stabilizes the actin cytoskeleton of neuronal processes by inducing n-cofilin phosphorylation at serine3. J. Neurosci. 29 (2009) 288–299. 10.1523/jneurosci.2934-08.2009.

[23] J.T. Rogers, I. Rusiana, J. Trotter, L. Zhao, E. Donaldson, D.T. Pak, L.W. Babus, M. Peters, J.L. Banko, P. Chavis, G.W. Rebeck, H.S. Hoe, & E.J. Weeber. Reelin supplementation enhances cognitive ability, synaptic plasticity, and dendritic spine density. Learn. Mem. 18 (2011) 558–564. 10.1101/lm.2153511.

[24] V.S. Caviness, Jr., & P. Rakic. Mechanisms of cortical development: a view from mutations in mice. Annu. Rev. Neurosci. 1 (1978) 297–326. 10.1146/annurev.ne.01.030178.001501.

[25] S. Hirotsune, T. Takahara, N. Sasaki, K. Hirose, A. Yoshiki, T. Ohashi, M. Kusakabe, Y. Murakami, M. Muramatsu, & S. Watanabe. The reeler gene encodes a protein with an EGF-like motif expressed by pioneer neurons. Nat. Genet. 10 (1995) 77–83. 10.1038/ng0595-77.

[26] V. de Bergeyck, K. Nakajima, C. Lambert de Rouvroit, B. Naerhuyzen, A.M. Goffinet, T. Miyata, M. Ogawa, & K. Mikoshiba. A truncated Reelin protein is produced but not secreted in the ‘Orleans’ reeler mutation (Reln[rl-Orl]). Brain Res. Mol. Brain Res. 50 (1997) 85–90. 10.1016/s0169-328x(97)00166-6.

[27] Y. Katsuyama & T. Terashima. Developmental anatomy of reeler mutant mouse. Dev. Growth Differ. 51 (2009) 271–286. 10.1111/j.1440-169X.2009.01102.x.

[28] S. Qiu, K.M. Korwek, A.R. Pratt-Davis, M. Peters, M.Y. Bergman, & E.J. Weeber. Cognitive disruption and altered hippocampus synaptic function in Reelin haploinsufficient mice. Neurobiol. Learn. Mem. 85 (2006) 228–242. 10.1016/j.nlm.2005.11.001.

[29] P. Tueting, E. Costa, Y. Dwivedi, A. Guidotti, F. Impagnatiello, R. Manev, & C. Pesold. The phenotypic characteristics of heterozygous reeler mouse. Neuroreport. 10 (1999) 1329–1334. 10.1097/00001756-199904260-00032.

[30] W.L. Salinger, P. Ladrow, & C. Wheeler. Behavioral phenotype of the reeler mutant mouse: effects of RELN gene dosage and social isolation. Behav. Neurosci. 117 (2003) 1257–1275. 10.1037/0735-7044.117.6.1257.

[31] W.S. Liu, C. Pesold, M.A. Rodriguez, G. Carboni, J. Auta, P. Lacor, J. Larson, B.G. Condie, A. Guidotti, & E. Costa. Down-regulation of dendritic spine and glutamic acid decarboxylase 67 expressions in the reelin haploinsufficient heterozygous reeler mouse. Proc. Natl. Acad. Sci. U. S. A. 98 (2001) 3477–3482. 10.1073/pnas.051614698.

[32] M. Poukkula, E. Kremneva, M. Serlachius, & P. Lappalainen. Actin-depolymerizing factor homology domain: A conserved fold performing diverse roles in cytoskeletal dynamics. Cytoskeleton. 68 (2011) 471–490. 10.1002/cm.20530.

[33] G.M. Beaudoin 3rd, S.-H. Lee, D. Singh, Y. Yuan, Y.G. Ng, L.F. Reichardt, & J. Arikkath. Culturing pyramidal neurons from the early postnatal mouse hippocampus and cortex. Nat. Protoc. 7 (2012) 1741–1754. 10.1038/nprot.2012.099.

[34] N. Yasui, T. Nogi, T. Kitao, Y. Nakano, M. Hattori, & J. Takagi. Structure of a receptor-binding fragment of reelin and mutational analysis reveal a recognition mechanism similar to endocytic receptors. Proc. Natl. Acad. Sci. U. S. A. 104 (2007) 9988–9993. 10.1073/pnas.0700438104.

[35] M. Sawahata, H. Asano, T. Nagai, N. Ito, T. Kohno, T. Nabeshima, M. Hattori, & K. Yamada. Microinjection of Reelin into the mPFC prevents MK-801-induced recognition memory impairment in mice. Pharmacol. Res. 173 (2021) 105832. 10.1016/j.phrs.2021.105832.

[36] J.H. Hanke, J.P. Gardner, R.L. Dow, P.S. Changelian, W.H. Brissette, E.J. Weringer, B.A. Pollok, & P.A. Connelly. Discovery of a novel, potent, and Src family-selective tyrosine kinase inhibitor. Study of Lck- and FynT-dependent T cell activation. J. Biol. Chem. 271 (1996) 695–701. 10.1074/jbc.271.2.695.

[37] H. Yang, L. Wang, C. Zang, Y. Wang, J. Shang, Z. Zhang, H. Liu, X. Bao, X. Wang, & D. Zhang. Src Inhibition Attenuates Neuroinflammation and Protects Dopaminergic Neurons in Parkinson’s Disease Models. Front. Neurosci. 14 (2020) 45. 10.3389/fnins.2020.00045.

[38] L. O’Donoghue & A. Smolenski. Analysis of protein phosphorylation using Phos-tag gels. J. Proteomics. 259 (2022) 104558. 10.1016/j.jprot.2022.104558.

[39] H. Ulrichs, I. Gaska, & S. Shekhar. Multicomponent regulation of actin barbed end assembly by twinfilin, formin and capping protein. Nat. Commun. 14 (2023) 3981. 10.1038/s41467-023-39655-3.

[40] K.R. Myers, Y.J. Fan, P. McConnell, J.A. Cooper, & J.Q. Zheng. Actin capping protein regulates postsynaptic spine development through CPI-motif interactions. Front. Mol. Neurosci. 15 (2022) 1020949. 10.3389/fnmol.2022.1020949.

[41] M. Hakala, H. Wioland, M. Tolonen, T. Kotila, A. Jegou, G. Romet-Lemonne, & P. Lappalainen. Twinfilin uncaps filament barbed ends to promote turnover of lamellipodial actin networks. Nat. Cell Biol. 23 (2021) 147–159. 10.1038/s41556-020-00629-y.

[42] L. A. Cingolani & Y. Goda. Actin in action: the interplay between the actin cytoskeleton and synaptic efficacy. Nat. Rev. Neurosci. 9 (2008) 344–356. 10.1038/nrn2373.

[43] I. Hlushchenko, P. Khanal, A. Abouelezz, V. O. Paavilainen, & P. Hotulainen. ASD-Associated De Novo Mutations in Five Actin Regulators Show Both Shared and Distinct Defects in Dendritic Spines and Inhibitory Synapses in Cultured Hippocampal Neurons. Front. Cell. Neurosci. 12 (2018) 217. 10.3389/fncel.2018.00217.

[44] R. Tanaka, J. Liao, K. Hada, D. Mori, T. Nagai, T. Matsuzaki, T. Nabeshima, K. Kaibuchi, N. Ozaki, H. Mizoguchi, & K. Yamada. Inhibition of Rho-kinase ameliorates decreased spine density in the medial prefrontal cortex and methamphetamine-induced cognitive dysfunction in mice carrying schizophrenia-associated mutations of the Arhgap10 gene. Pharmacol. Res. 187 (2023) 106589. 10.1016/j.phrs.2022.106589.

[45] J. Liao, G. Dong, B. Wulaer, M. Sawahata, H. Mizoguchi, D. Mori, N. Ozaki, T. Nabeshima, T. Nagai, & K. Yamada. Mice with exonic RELN deletion identified from a patient with schizophrenia have impaired visual discrimination learning and reversal learning in touchscreen operant tasks. Behav. Brain Res. 416 (2022) 113569. 10.1016/j.bbr.2021.113569.

[46] A.E. Horner, C.J. Heath, M. Hvoslef-Eide, B.A. Kent, C.H. Kim, S.R. Nilsson, J. Alsiö, C.A. Oomen, A. Holmes, L.M. Saksida, & T.J. Bussey. The touchscreen operant platform for testing learning and memory in rats and mice. Nat. Protoc. 8 (2013) 1961–1984. 10.1038/nprot.2013.122.

[47] J.L. Brigman, R.A. Daut, T. Wright, O. Gunduz-Cinar, C. Graybeal, M.I. Davis, Z. Jiang, L.M. Saksida, S. Jinde, M. Pease, T.J. Bussey, D.M. Lovinger, K. Nakazawa, & A. Holmes. GluN2B in corticostriatal circuits governs choice learning and choice shifting. Nat. Neurosci. 16 (2013) 1101–1110. 10.1038/nn.3457.

[48] S.A. Bonini, A. Mastinu, G. Maccarinelli, S. Mitola, M. Premoli, L.R. La Rosa, G. Ferrari-Toninelli, M. Grilli, & M. Memo. Cortical Structure Alterations and Social Behavior Impairment in p50-Deficient Mice. Cereb. Cortex. 26 (2016) 2832–2849. 10.1093/cercor/bhw037.

[49] H. Tabata, D. Mori, T. Matsuki, K. Yoshizaki, M. Asai, A. Nakayama, N. Ozaki, & K.I. Nagata. Histological Analysis of a Mouse Model of the 22q11.2 Microdeletion Syndrome. Biomolecules. 13 (2023) 763. 10.3390/biom13050763.

[50] M. Sawahata, D. Mori, Y. Arioka, H. Kubo, I. Kushima, K. Kitagawa, A. Sobue, E. Shishido, M. Sekiguchi, A. Kodama, R. Ikeda, B. Aleksic, H. Kimura, K. Ishizuka, T. Nagai, K. Kaibuchi, T. Nabeshima, K. Yamada, & N. Ozaki. Generation and analysis of novel Reln-deleted mouse model corresponding to exonic Reln deletion in schizophrenia. Psychiatry Clin. Neurosci. 74 (2020) 318–327. 10.1111/pcn.12993.

[51] P. Xu, A. Chen, Y. Li, X. Xing, & H. Lu. Medial prefrontal cortex in neurological diseases. Physiol. Genomics. 51 (2019) 432–442. 10.1152/physiolgenomics.00006.2019.

[52] J.A. Sexton, T. Potchernikov, J.P. Bibeau, G. Casanova-Sepúlveda, W. Cao, H.J. Lou, T.J. Boggon, E.M. De La Cruz, & B.E. Turk. Distinct functional constraints driving conservation of the cofilin N-terminal regulatory tail. Nat. Commun. 15 (2024) 1426. 10.1038/s41467-024-45878-9.

[53] Y. Arioka, E. Shishido, H. Kubo, I. Kushima, A. Yoshimi, H. Kimura, K. Ishizuka, B. Aleksic, T. Maeda, M. Ishikawa, N. Kuzumaki, H. Okano, D. Mori, & N. Ozaki. Single-cell trajectory analysis of human homogenous neurons carrying a rare RELN variant. Transl. Psychiatry. 8 (2018) 129. 10.1038/s41398-018-0177-8.

[54] J.S. da Silva, & C.G. Dotti. Breaking the neuronal sphere: regulation of the actin cytoskeleton in neuritogenesis. Nat. Rev. Neurosci. 3 (2002) 694–704. 10.1038/nrn918.

[55] X. Chai, L. Fan, H. Shao, X. Lu, W. Zhang, J. Li, J. Wang, S. Chen, M. Frotscher, & S. Zhao. Reelin Induces Branching of Neurons and Radial Glial Cells during Corticogenesis. Cereb. Cortex. 25 (2015) 3640–3653. 10.1093/cercor/bhu216.

[56] S. Qiu & E.J. Weeber. Reelin signaling facilitates maturation of CA1 glutamatergic synapses. J. Neurophysiol. 97 (2007) 2312–2321. 10.1152/jn.00869.2006.

[57] Y. Chen, U. Beffert, M. Ertunc, T.S. Tang, E.T. Kavalali, I. Bezprozvanny, & J. Herz. Reelin modulates NMDA receptor activity in cortical neurons. J. Neurosci. 25 (2005) 8209–8216. 10.1523/jneurosci.1951-05.2005.

[58] M.K. Vartiainen, E.M. Sarkkinen, T. Matilainen, M. Salminen, & P. Lappalainen. Mammals have two twinfilin isoforms whose subcellular localizations and tissue distributions are differentially regulated. J. Biol. Chem. 278 (2003) 34347–34355. 10.1074/jbc.M303642200.

[59] M. Vartiainen, P.J. Ojala, P. Auvinen, J. Peränen, & P. Lappalainen. Mouse A6/twinfilin is an actin monomer-binding protein that localizes to the regions of rapid actin dynamics. Mol. Cell. Biol. 20 (2000) 1772–1783. 10.1128/mcb.20.5.1772-1783.2000.

[60] B.L. Goode, D.G. Drubin, & P. Lappalainen. Regulation of the cortical actin cytoskeleton in budding yeast by twinfilin, a ubiquitous actin monomer-sequestering protein. J. Cell Biol. 142 (1998) 723–733. 10.1083/jcb.142.3.723.

[61] D. Wang, L. Zhang, G. Zhao, G. Wahlström, T.I. Heino, J. Chen, & Y.Q. Zhang. Drosophila twinfilin is required for cell migration and synaptic endocytosis. J. Cell Sci. 123 (2010) 1546–1556. 10.1242/jcs.060251.

[62] S. Shekhar, G.J. Hoeprich, J. Gelles, & B.L. Goode. Twinfilin bypasses assembly conditions and actin filament aging to drive barbed end depolymerization. J. Cell Biol. 220 (2021) e202006022. 10.1083/jcb.202006022.

[63] H. Hering & M. Sheng. Dendritic spines: structure, dynamics and regulation. Nat. Rev. Neurosci. 2 (2001) 880–888. 10.1038/35104061.

[64] L. Luo. Actin cytoskeleton regulation in neuronal morphogenesis and structural plasticity. Annu. Rev. Cell Dev. Biol. 18 (2002) 601–635. 10.1146/annurev.cellbio.18.031802.150501.

[65] C. Dillon & Y. Goda. The actin cytoskeleton: integrating form and function at the synapse. Annu. Rev. Neurosci. 28 (2005) 25–55. 10.1146/annurev.neuro.28.061604.135757.

[66] A.B. Johnston, D.M. Hilton, P. McConnell, B. Johnson, M.T. Harris, A. Simone, G.K. Amarasinghe, J.A. Cooper, & B.L. Goode. A novel mode of capping protein-regulation by twinfilin. Elife. 7 (2018) e41313. 10.7554/eLife.41313.

[67] T.J. Bussey, A. Holmes, L. Lyon, A.C. Mar, K.A. McAllister, J. Nithianantharajah, C.A. Oomen, & L.M. Saksida. New translational assays for preclinical modelling of cognition in schizophrenia: the touchscreen testing method for mice and rats. Neuropharmacology. 62 (2012) 1191–1203. 10.1016/j.neuropharm.2011.04.011.

[68] M.P. Boyle, A. Bernard, C.L. Thompson, L. Ng, A. Boe, M. Mortrud, M.J. Hawrylycz, A.R. Jones, R.F. Hevner, & E.S. Lein. Cell-type-specific consequences of Reelin deficiency in the mouse neocortex, hippocampus, and amygdala. J. Comp. Neurol. 519 (2011) 2061–2089. 10.1002/cne.22655.

[69] K. Sakai, H. Shoji, T. Kohno, T. Miyakawa, & M. Hattori. Mice that lack the C-terminal region of Reelin exhibit behavioral abnormalities related to neuropsychiatric disorders. Sci. Rep. 6 (2016) 28636. 10.1038/srep28636.

[70] R. Lalonde, K. Hayzoun, M. Derer, J. Mariani, & C. Strazielle. Neurobehavioral evaluation of Reln-rl-orl mutant mice and correlations with cytochrome oxidase activity. Neurosci. Res. 49 (2004) 297–305. 10.1016/j.neures.2004.03.012.

[71] S. Niu, A. Renfro, C.C. Quattrocchi, M. Sheldon, & G. D’Arcangelo. Reelin promotes hippocampal dendrite development through the VLDLR/ApoER2-Dab1 pathway. Neuron. 41 (2004) 71–84. 10.1016/s0896-6273(03)00819-5.

[72] M.C. Pinto Lord & V.S. Caviness Jr. Determinants of cell shape and orientation: a comparative Golgi analysis of cell-axon interrelationships in the developing neocortex of normal and reeler mice. J. Comp. Neurol. 187 (1979) 49–69. 10.1002/cne.901870104.

[73] A.M. Goffinet & G. Lyon. Early histogenesis in the mouse cerebral cortex: a Golgi study. Neurosci. Lett. 14 (1979) 61–66. 10.1016/0304-3940(79)95344-8.

[74] C.M. Teixeira, M.M. Kron, N. Masachs, H. Zhang, D.C. Lagace, A. Martinez, I. Reillo, X. Duan, C. Bosch, L. Pujadas, L. Brunso, H. Song, A.J. Eisch, V. Borrell, B.W. Howell, J.M. Parent, & E. Soriano. Cell-autonomous inactivation of the reelin pathway impairs adult neurogenesis in the hippocampus. J. Neurosci. 32 (2012) 12051–12065. 10.1523/jneurosci.1857-12.2012.

[75] M. Sinagra, D. Verrier, D. Frankova, K.M. Korwek, J. Blahos, E.J. Weeber, O.J. Manzoni, & P. Chavis. Reelin, very-low-density lipoprotein receptor, and apolipoprotein E receptor 2 control somatic NMDA receptor composition during hippocampal maturation in vitro. J. Neurosci. 25 (2005) 6127–6136. 10.1523/jneurosci.1757-05.2005.

[76] Y. Tsuneura, M. Sawahata, N. Itoh, R. Miyajima, D. Mori, T. Kohno, M. Hattori, A. Sobue, T. Nagai, H. Mizoguchi, T. Nabeshima, N. Ozaki, & K. Yamada. Analysis of Reelin signaling and neurodevelopmental trajectory in primary cultured cortical neurons with RELN deletion identified in schizophrenia. Neurochem. Int. 144 (2021) 104954. 10.1016/j.neuint.2020.104954.

[77] B.S. Chang, F. Duzcan, S. Kim, M. Cinbis, A. Aggarwal, K.A. Apse, O. Ozdel, M. Atmaca, S. Zencir, H. Bagci, & C.A. Walsh. The role of RELN in lissencephaly and neuropsychiatric disease. Am. J. Med. Genet. B Neuropsychiatr. Genet. 144b (2007) 58–63. 10.1002/ajmg.b.30392.

[78] D.A. Skaar, Y. Shao, J.L. Haines, J.E. Stenger, J. Jaworski, E.R. Martin, G.R. DeLong, J.H. Moore, J.L. McCauley, J.S. Sutcliffe, A.E. Ashley-Koch, M.L. Cuccaro, S.E. Folstein, J.R. Gilbert, & M.A. Pericak-Vance. Analysis of the RELN gene as a genetic risk factor for autism. Mol. Psychiatry. 10 (2005) 563–571. 10.1038/sj.mp.4001614.

[79] D.R. Grayson, X. Jia, Y. Chen, R.P. Sharma, C.P. Mitchell, A. Guidotti, & E. Costa. Reelin promoter hypermethylation in schizophrenia. Proc. Natl. Acad. Sci. U. S. A. 102 (2005) 9341–9346. 10.1073/pnas.0503736102.

[80] R.M. Nabil Fikri, A.T. Norlelawati, A.R. Nour El-Huda, M.N. Hanisah, A. Kartini, K. Norsidah, & A. Nor Zamzila. Reelin (RELN) DNA methylation in the peripheral blood of schizophrenia. J. Psychiatr. Res. 88 (2017) 28–37. 10.1016/j.jpsychires.2016.12.020.

[81] J. Jackson, E. Jambrina, J. Li, H. Marston, F. Menzies, K. Phillips, & G. Gilmour. Targeting the Synapse in Alzheimer’s Disease. Front. Neurosci. 13 (2019) 735. 10.3389/fnins.2019.00735.

[82] Y. Arioka, A. Hirata, I. Kushima, B. Aleksic, D. Mori, & N. Ozaki. Characterization of a schizophrenia patient with a rare RELN deletion by combining genomic and patient-derived cell analyses. Schizophr. Res. 216 (2020) 511–515. 10.1016/j.schres.2019.10.038.

[83] T. Aida, K. Chiyo, T. Usami, H. Ishikubo, R. Imahashi, Y. Wada, K.F. Tanaka, T. Sakuma, T. Yamamoto, & K. Tanaka. Cloning-free CRISPR/Cas system facilitates functional cassette knock-in in mice. Genome Biol. 16 (2015) 87. 10.1186/s13059-015-0653-x

[84] D. Mori, R. Ikeda, M. Sawahata, S. Yamaguchi, A. Kodama, T. Hirao, Y. Arioka, H. Okumura, C. Inami, T. Suzuki, Y. Hayashi, H. Kato, Y. Nawa, S. Miyata, H. Kimura, I. Kushima, B. Aleksic, H. Mizoguchi, T. Nagai, T. Nakazawa, R. Hashimoto, K. Kaibuchi, K. Kume, K. Yamada, & N. Ozaki. Phenotypes for general behavior, activity, and body temperature in 3q29 deletion model mice. Transl. Psychiatry. 14 (2024) 138. 10.1038/s41398-023-02679-w.

[85] D. Mori, C. Inami, R. Ikeda, M. Sawahata, S. Urata, S.T. Yamaguchi, Y. Kobayashi, K. Fujita, Y. Arioka, H. Okumura, I. Kushima, A. Kodama, T. Suzuki, T. Hirao, A. Yoshimi, A. Sobue, T. Ito, Y. Noda, H. Mizoguchi, T. Nagai, K. Kaibuchi, S. Okabe, K. Nishiguchi, K. Kume, K. Yamada, & N. Ozaki. Mice with deficiency in Pcdh15, a gene associated with bipolar disorders, exhibit significantly elevated diurnal amplitudes of locomotion and body temperature. Transl. Psychiatry. 14 (2024) 216. 10.1038/s41398-024-02952-6.

[86] M. Sekiguchi, A. Sobue, I. Kushima, C. Wang, Y. Arioka, H. Kato, A. Kodama, H. Kubo, N. Ito, M. Sawahata, K. Hada, R. Ikeda, M. Shinno, C. Mizukoshi, K. Tsujimura, A. Yoshimi, K. Ishizuka, Y. Takasaki, H. Kimura, J. Xing, Y. Yu, M. Yamamoto, T. Okada, E. Shishido, T. Inada, M. Nakatochi, T. Takano, K. Kuroda, M. Amano, B. Aleksic, T. Yamomoto, T. Sakuma, T. Aida, K. Tanaka, R. Hashimoto, M. Arai, M. Ikeda, N. Iwata, T. Shimamura, T. Nagai, T. Nabeshima, K. Kaibuchi, K. Yamada, D. Mori, & N. Ozaki. ARHGAP10, which encodes Rho GTPase-activating protein 10, is a novel gene for schizophrenia risk. Transl. Psychiatry. 10 (2020) 247. 10.1038/s41398-020-00917-z.

[87] S. Yamada, T. Nagai, T. Nakai, D. Ibi, A. Nakajima, & K. Yamada. Matrix metalloproteinase-3 is a possible mediator of neurodevelopmental impairment due to polyI:C-induced innate immune activation of astrocytes. Brain Behav. Immun. 38 (2014) 272–282. 10.1016/j.bbi.2014.02.014.

[88] T. Nakai, T. Nagai, M. Tanaka, N. Itoh, N. Asai, A. Enomoto, M. Asai, S. Yamada, A.B. Saifullah, M. Sokabe, M. Takahashi, & K. Yamada. Girdin phosphorylation is crucial for synaptic plasticity and memory: a potential role in the interaction of BDNF/TrkB/Akt signaling with NMDA receptor. J. Neurosci. 34 (2014) 14995–15008. 10.1523/jneurosci.2228-14.2014.

[89] K.J. Livak & T.D. Schmittgen. Analysis of relative gene expression data using real-time quantitative PCR and the 2(-Delta Delta C(T)) Method. Methods. 25 (2001) 402–408. 10.1006/meth.2001.1262.

[90] A. Sobue, N. Ito, T. Nagai, W. Shan, K. Hada, A. Nakajima, Y. Murakami, A. Mouri, Y. Yamamoto, T. Nabeshima, K. Saito, & K. Yamada. Astroglial major histocompatibility complex class I following immune activation leads to behavioral and neuropathological changes. Glia. 66 (2018) 1034–1052. 10.1002/glia.23299.

[91] J. Liao, G. Dong, W. Zhu, B. Wulaer, H. Mizoguchi, M. Sawahata, Y. Liu, K. Kaibuchi, N. Ozaki, T. Nabeshima, T. Nagai, & K. Yamada. Rho kinase inhibitors ameliorate cognitive impairment in a male mouse model of methamphetamine-induced schizophrenia. Pharmacol. Res. 194 (2023) 106838. 10.1016/j.phrs.2023.106838.

[92] D. Ibi, T. Nagai, H. Koike, Y. Kitahara, H. Mizoguchi, M. Niwa, H. Jaaro-Peled, A. Nitta, Y. Yoneda, T. Nabeshima, A. Sawa, & K. Yamada. Combined effect of neonatal immune activation and mutant DISC1 on phenotypic changes in adulthood. Behav. Brain Res. 206 (2010) 32–37. 10.1016/j.bbr.2009.08.027.

[93] H. Kubota, K. Kunisawa, B. Wulaer, M. Hasegawa, H. Kurahashi, T. Sakata, H. Tezuka, M. Kugita, S. Nagao, T. Nagai, T. Furuyashiki, S. Narumiya, K. Saito, T. Nabeshima, & A. Mouri. High salt induces cognitive impairment via the interaction of the angiotensin II-AT(1) and prostaglandin E2-EP(1) systems. Br. J. Pharmacol. 180 (2023) 2393–2411. 10.1111/bph.16093.

[94] B. Wulaer, T. Nagai, A. Sobue, N. Itoh, K. Kuroda, K. Kaibuchi, T. Nabeshima, & K. Yamada. Repetitive and compulsive-like behaviors lead to cognitive dysfunction in Disc1(Δ2-3/Δ2-3) mice. Genes Brain Behav. 17 (2018) e12478. 10.1111/gbb.12478.

[95] Y. Hayashi, T. Nagai, A. Sobue, N. Itoh, K. Kuroda, K. Kaibuchi, T. Nabeshima, & K. Yamada. Analysis of human neuronal cells carrying ASTN2 deletion associated with psychiatric disorders. Transl. Psychiatry. 14 (2024) 236. 10.1038/s41398-024-02962-4.

[96] Y. Arioka, E. Shishido, I. Kushima, T. Suzuki, R. Saito, A. Aiba, D. Mori, & N. Ozaki. Chromosome 22q11.2 deletion causes PERK-dependent vulnerability in dopaminergic neurons. EBioMedicine. 63 (2021) 103138. 10.1016/j.ebiom.2020.103138.

[97] G. Jacquemet, I. Paatero, A.F. Carisey, A. Padzik, J.S. Orange, H. Hamidi, & J. Ivaska. FiloQuant reveals increased filopodia density during breast cancer progression. J. Cell Biol. 216 (2017) 3387–3403. 10.1083/jcb.201704045.

